# Single-cell phenomics through integrated imaging and molecular profiling

**DOI:** 10.1101/2025.11.28.690954

**Authors:** Johannes Bues, Joern Pezoldt, Camille Lucie Lambert, Benjamin David Hale, Elisa Bugani, Ramon Vinas Torne, Timothee Ferrari, Nadia Grenningloh, Vincent Gardeux, Shuo Wen, Caroline Wandinger, Maximilian Kohnen, Romina Augustin, Katharina Eckstein, Assia Ouanaya, Jillian Love, Sarthak Saha, Amirhossein Saba, Aviv Huttner, Maria Vittoria Impagliazzo, Jose Antonio Vasquez Porto Viso, Angel de Jesus Corria Osorio, Demetri Psaltis, Wouter Karthaus, Berend Snijder, Maria Brbic, Bart Deplancke

## Abstract

Single-cell technologies such as transcriptomics, microscopy, and flow cytometry have revolutionized the study of cellular identity and function. While each of these technologies is powerful on its own, their full potential lies in their integration, enabling multimodal profiling of the same cell and revealing how distinct modalities influence one another. Here, we introduce IRIS (Interconnected Robotic Imaging and Single cell transcriptomics), a deterministic single-cell platform technology that seamlessly couples high-resolution microscopy with droplet-based single-cell RNA sequencing. IRIS enables precise cell positioning, multimode imaging across brightfield and fluorescent channels, and subsequent molecular capture from the same cell, directly linking high-resolution morphological features to matched transcriptomes. We validate IRIS by recovering cell cycle progression states and transcriptional programmes associated with canonical morphologies and demonstrate its discovery power by molecularly resolving two nuclear-ER architectures within naïve CD8^+^ T cells, each defined by distinct gene expression profiles and functional markers. IRIS establishes an integrative single-cell phenomics framework, opening new avenues for dissecting how cellular form relates to molecular state and function.

## Introduction

Single-cell omics technologies have transformed our ability to dissect heterogeneity by enabling high-resolution molecular profiling of individual cells^1,2,3^. Yet, molecular profiles alone often fall short of explaining how cells behave, interact, or respond to their environment without a direct link to phenotypic properties. These properties, such as dynamic behaviours and structural properties, are increasingly recognized as essential for interpreting cellular identity and state^4,5^. Modern biology is therefore shifting toward a broader framework of ‘single-cell phenomics’, in which molecular measurements are paired with diverse phenotypic readouts to better resolve cell identity and function^6^.

On the phenotypic side, conventional single-cell imaging and flow-imaging systems capture rich morphological detail, including the spatial organization of organelles, membranes, and cytoskeletal elements. These phenotypic features are highly informative as they provide an architectural blueprint of cell function^7^, which holds both fundamental value for understanding cellular behaviour and direct clinical relevance: in haematology, for example, cell size and the shape of blast cells are central to diagnosing and classifying acute myeloid leukemia^8^, while morphological characteristics of tumour cells are a predictor of metastatic potential^9^. However, these imaging approaches lack the ability to recover molecular profiles from the same imaged cells^10,11^, making it difficult to interpret the functional relevance of the detected morphological features. Advances such as image-based cell sorting^12,13,10,14^ allow the isolation of cells by morphology for plate-based molecular profiling^10^. However, because these approaches rely on computationally reconstructed images rather than high-resolution microscopy, they effectively capture coarse cell properties such as size, granularity or refractive index, but are limited in resolving more fine-grained, morphological, or organelle-level features accessible with conventional microscopy^15,16^. Additionally, their reliance on plate-based profiling constrains molecular measurements due to the method’s labour-intensive, costly, and poorly scalable nature.

On the molecular side, multimodal strategies have broadened the scope of single-cell transcriptomics (scRNA-seq) by integrating complementary biological layers. For example, CITE-seq couples surface protein epitopes with transcriptomes^17^, Perturb-seq^18^ and sci-Plex^19^ connect transcriptional changes to genetic or chemical perturbations, PERISCOPE^20^ integrates high-dimensional imaging-based phenotyping with pooled CRISPR perturbations to link morphological changes to specific gene perturbations, and Live-seq links upstream molecular states to downstream cellular behaviour through transcriptomic profiling of still-viable cells^21^. Likewise, spatial omics defines molecular states within their native microenvironments, revealing how spatial organization and cell–cell interactions shape fate and function^22,23^. While these multimodal extensions enrich molecular readouts, they do not link dense transcriptomic data to high-resolution, single-cell morphological features. Some of the earliest scRNA-seq platforms did in fact offer a first step toward this integration by virtue of their design. Specifically, by utilizing hydrodynamic traps (Fluidigm C1^24^) or microwells (Takara/Wafergen iCell8^25^), these methods achieved deterministic cell capture, ensuring that each molecular profile could be traced back to a specific imaged cell. However, these tools offered only low-quality imaging (Takara/Wafergen iCell8^25^) or limited transcriptome sensitivity (Scope-Seq^26^), and throughput was constrained by the finite number of capture sites.

Subsequent scRNA-seq technologies overcame throughput limitations by introducing stochastic barcoding, encapsulating cells with barcoded beads in microwells^27^ or nanoliter droplets^28,29^. This major advance enabled profiling of thousands to millions of cells in a single experiment but at the expense of determinism, since random barcodes cannot be linked back to specific (e.g., imaged) cells. A recent approach partially addressed this by performing spatial transcriptomics on thousands of ‘slide-stamped’, fixed cells^30^. While this approach comes with trade-offs, including limited transcriptome coverage and the inability to perform live-cell staining or handling, it conceptually opens the path toward directly pairing molecular readouts with morphological features. Doing so would provide complementary context, as each modality captures distinct and only partially overlapping dimensions of cell identity. Such integration may also enable machine learning–driven inference between modalities, where phenotypic features inform molecular interpretation and vice versa, ultimately advancing a more predictive understanding of cell function. In sum, single-cell imaging and sequencing have each yielded transformative insights into cellular heterogeneity and behaviour, but a technology capable of combining high-resolution morphological profiling with full transcriptomic coverage in the same cell, at scale, within a unified workflow has so far remained out of reach^6^.

Here, we present IRIS (Interconnected Robotic Imaging and Sequencing), the first platform to unite deterministic microfluidics, machine vision-guided high-resolution imaging, and dual-indexed barcoding. IRIS achieves precise one-to-one pairing of submicron-resolved morphological features with transcriptomes, while retaining scalability and sensitivity. This positions IRIS to fill a longstanding gap in multimodal profiling, delivering genome-wide gene expression profiles together with high-resolution morphology to resolve dynamic cellular states. We demonstrate the power of this approach by benchmarking morphological features across the cell cycle and uncovering molecular programs associated with distinct morpho-architectural classes of naïve CD8+ T cells, establishing IRIS as a foundation for integrated single-cell phenomics at scale.

## Results

### Deterministic microfluidic handling of single cells enables native multiplexing

To bridge phenotypic and molecular profiling, we established a technology based on four core properties: (1) stopped-flow cell handling to allow extended acquisition of high-resolution phenotypic data, (2) molecular manipulation within scalable nanoliter droplets, (3) use of in-suspension biochemical reactants for maximum omics protocol flexibility, and (4) full process and barcode determinism for reliably linking phenotype (image) and transcriptome.

Our solution, overviewed in **Fig. 1**, is a hybrid platform that leverages microfluidics for real-time single-cell manipulation, multiwell plates for deterministic cell tracking, and nanoliter droplet biochemistry for scalable processing. In brief, individual cells are optically detected on-chip, stopped via machine vision-triggered solenoid-actuated Quake valves^31^ (**Fig. 1**, **Fig. 2A**), and precisely positioned in a channel segment by over-pressurizing the leak-flow Quake valve for high-resolution imaging (**Fig. 2A, Supplementary Fig. 2A, Video 1**). Each cell is then co-injected into a droplet by dispensing the reaction mix from an additional inlet, producing a 5 nL plug composed of equal volumes of reaction mix and cell buffer. This plug is then sheared by oil to form an individual droplet. The reaction mix contains a reverse transcription (RT) poly-dT primer including a predefined Cell Code (CC) and a unique molecular identifier (UMI) (**Fig. 2B**). Each droplet is subsequently deposited into a designated well of a 96-well plate (system depicted in **Supplementary Fig. 2B**), establishing a one-to-one correspondence between each captured cell and its well location, thereby ensuring complete traceability.

**Figure 1.**
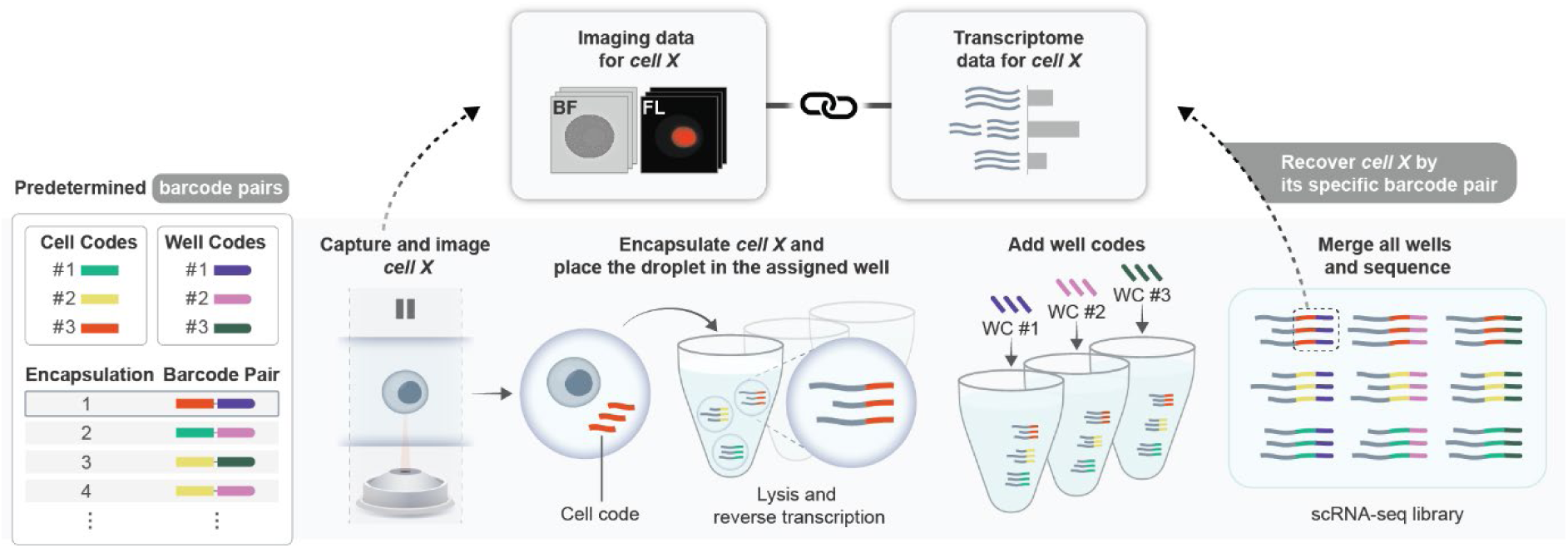
Overview of Integrated Robotic Imaging and Sequencing, IRIS. The IRIS platform enables one-to-one linkage between cell images and transcriptomic profiles through a deterministic, microfluidic workflow. Individual cells are optically detected, stopped, and imaged with brightfield (BF) and fluorescence (FL) microscopy. Each imaged cell is then encapsulated in a nanoliter droplet with a unique Cell Code (CC) and deposited into a predefined well of a 96-well plate. This results in a unique mapping of each cell and its CC to a specific well, preserving cell identity. Following droplet deposition, cells are lysed and mRNA is reverse transcribed in all droplets within each well. Subsequently, a second barcode, the Well Code (WC), is added via overhang PCR. This dual-indexing strategy (CC + WC) allows pooled sequencing of all wells across the plate, enabling precise reconstruction of transcriptomes for each imaged cell. In this way, the IRIS platform creates a barcode pair that uniquely links each cell’s phenotype (image) to its gene expression profile. Multiple barcode-labelled droplets can be processed in a single well using this ‘droplet consortia’ method, maximizing scalability without compromising traceability. The final output is a merged single-cell RNA-seq library, where each transcriptome is linked to its corresponding imaging data. This integrated approach provides scalable phenotypic and molecular profiling at single-cell resolution.

**Fig. 2.**
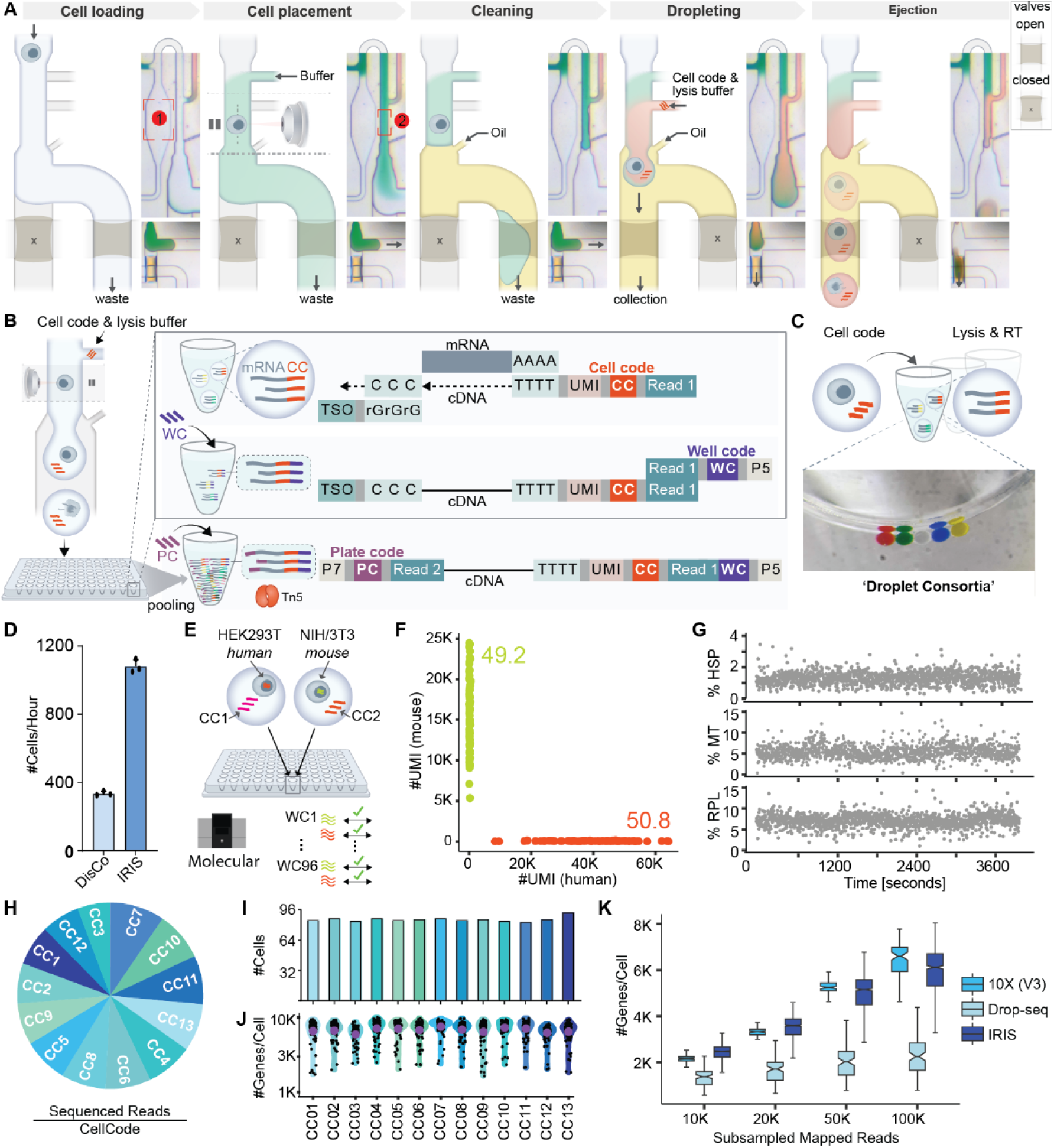
‘Droplet consortia’ enable native multiplexing. **(A)** Depicts the deterministic single-cell encapsulation process using Quake valve-based leak-flow control. *From left to right*: Cells are first loaded into the microfluidic chip via the main loading channel. They are decelerated in a widening region (1) and subsequently positioned in the imaging zone (2) through an injection channel (green buffer), where they will be stopped for imaging. The downstream channel is then flushed with oil, and the Cell Code (CC) and lysis buffer (red buffer) are injected, forming a bud within the oil phase that contains the cell. The resulting droplet is sheared and directed into the collection channel by closing the waste valve, where it is subsequently deposited into wells. **(B)** IRIS molecular workflow for deterministic per cell dual-indexed barcoding. Cells are stopped and encapsulated in droplets containing RT mix with a predefined CellCode (CC). Sequential ‘barcode sampling’ introduces a distinct CC for each set of 96 cells, with droplets deposited into assigned wells to form ‘droplet consortia’ with each droplet containing a different CC. A WellCode (WC) is added post-RT during PCR, producing a unique CC+WC combination (dual-index barcoding) that links each cell to its capture event (and imaging). The Plate Code (PC) is added during Tn5-based sequencing library preparation. **(C)** Example consortium of droplets loaded with dyes for visualization in a well of a 96-well plate. **(D)** Processing throughput of DisCo vs. IRIS. **(E)** Species-mixing design to assess droplet integrity: human HEK293T and mouse NIH/3T3 cells, each with a designated CC, were combined in the same well. Species separation was verified using scRNA-seq by mapping transcriptomic reads to a merged human-mouse reference genome. **(F)** Barnyard plot showing species separation from (E). **(G)** Continuous processing of HEK293T cells showing stable mitochondrial (MT), ribosomal protein (RPL), and heat-shock protein (HSP) RNA percentages over time. **(H-J)** Native multiplexing of 13 cells per well by iterative CC introduction. **(H)** Read proportions per CC. **(I)** Number of recovered cells (>3000 UMI/cell) per CC. **(J)** UMIs per cell for each CC RT-primer. **(K)** Genes per cell vs. down-sampled mapped reads of IRIS, Drop-seq (on DisCo) and 10x Genomics 3’ V3.

Importantly, droplets isolate the biochemical reaction, enabling multiple cells to be processed per well (**Fig. 2C**). To resolve these cells, however, each droplet must contain a distinguishable barcode, hence the CC. To achieve this, we iteratively process 96 cells per RT solution; each set of 96 cells is tagged with a different CC, and then we replace the RT mix to create a group of droplets with distinct and defined contents within each well, which we term ‘droplet consortia’ (**Fig. 2C**). Following RT, a second index, i.e. a Well Code (WC), is introduced via overhang PCR. This dual-indexing (CC+WC) strategy allows reconstruction of each cell’s transcriptome after pooling a 96-well plate and sequencing, while assuring robust, end-to-end linkage between image acquisition and molecular profiling (**Fig. 2B**). The optimized process reached a throughput of 1,150 cells/h (**Fig. 2D**), approximately three times faster than our previous deterministic co-encapsulation, low cell input scRNA-seq platform DisCo^31^, which provided the conceptual foundation for IRIS.

Together, this strategy overcomes longstanding technical limitations in linking morphology to molecular state at scale^6^. It provides an integrated, deterministic workflow in which phenotypic imaging and transcriptomic profiling are coupled at single-cell resolution.

To implement IRIS at scale, while ensuring molecular fidelity, we optimized key workflow components, including reaction biochemistry, barcode integrity, and continuous cell processing. These steps were essential to maximize cDNA yield, ensure accurate cell-to-barcode linkage, and preserve transcriptomic quality during continuous operation. RT efficiency^28,32^ was maximized by tuning reaction temperature, RT enzyme concentration, dCTP levels, and detergent concentration (**Supplementary Fig. 2C**). Library purity (e.g. removing residual WC primers) was improved through a two-step column-based cleanup and bead-based size selection (**Supplementary Fig. 2D-F**).

Barcode switching and RT solution exchange fidelity were tested by sequentially processing human HEK293T and mouse NIH/3T3 cells with distinct cell barcodes (CC1, CC2), placing one cell of each species in each well (**Fig. 2E**). All cells mapped exclusively to their expected species (98.8% of the transcripts for mouse and 99.8% for human on a per-cell basis), confirming effective RT-solution exchange and stable droplets (**Fig. 2F**). Unlike non-deterministic microfluidic systems^29,28,32^, IRIS-based molecular profiling operates continuously. By implementing a cooled cell mixer, we were able to process cells for over one hour without detectable stress induction, as shown by consistent proportions of mitochondrial, heat-shock protein, and ribosomal protein RNA fractions over time (**Fig. 2G**). Transcriptome quality (UMIs, gene counts) also remained constant over time with some variation in capture efficiency across RT primers (mean 23,000 ± 3,500 UMIs and 6,100 ± 350 genes; **Supplementary Fig. 2G-H**) and nearly consistent UMI yields across wells (**Supplementary Fig. 2I**).

To test IRIS’s native multiplexing capability, we profiled human HEK293T cells by generating ‘droplet consortia’ composed of 13 distinct RT primers, each with a unique CC. All CC primers contributed uniformly to the total read distribution (**Fig. 2H**), yielding on average 86.2 ± 2.5 cells per CC and 1,118 cells per 96-well plate (**Fig. 2I**). Sensitivity was maintained across primers, and, UMI and gene detection remained robust across RT primers, with 34,100 ± 4,300 UMIs and 7,040 ± 450 genes per cell, respectively (**Fig. 2J, Supplementary Fig. 2J**). Compared to existing droplet-based platforms, IRIS’s ‘droplet consortia’-based molecular profiling system outperformed Drop-seq by 2.6-fold in gene detection sensitivity and reached 92% of the performance of 10X Genomics v3 chemistry at 100,000 mapped reads per HEK293T cell (**Fig. 2K, Supplementary Fig. 2K-L**). This was achieved at a material cost of approximately $0.16 per cell, underscoring the scalability and affordability of the approach (**Supplementary Fig. 2M**).

Taken together, IRIS represents, to our knowledge, the first high-throughput platform combining deterministic cell handling, imaging, flexible nanoliter-scale droplet biochemistry and precise dual-indexed molecular tracking, enabling scalable, cost-effective integration of cell morphology and transcriptomics.

### Deterministic barcoding links imaging with transcriptomics at the single-cell level

To pair imaging with transcriptomic profiling, IRIS uses the machine vision-guided stopping and encapsulation strategy described above (**Fig. 2A**), in which decelerating cells are identified using real-time image subtraction and object detection algorithms, and then precisely positioned into a stopping zone (**Supplementary Fig. 3A**) for high-resolution imaging (∼500 nm lateral resolution using a 40X, NA 0.9 objective) before droplet encapsulation (**Methods**).

The microscope, equipped with a piezoelectric stage, acquires one brightfield and four epi-fluorescence channels (DAPI, FITC, TRITC, Cy5 or similar) per focal plane (**Fig. 3A**). Focal planes are acquired in stacked fashion across the channel height with adjustable inter-plane distances, ensuring optimal focus for each cell and enabling visualization of the 3D distribution of intracellular structures (**Fig. 3B-C**). Cropped cell images are extracted from the raw images with a YOLO-based object detection model that simultaneously identifies the cell’s location and the optimal focal plane from the brightfield data (**Fig. 3C** and **Supplementary Fig. 3B**, **Methods**). Additionally, we used a customized DeepLabV3 network to compute a segmentation mask from the in-focus brightfield image (**Supplementary Fig. 3C-D**, **Methods**), used for quality control, filtering, and intensity measurements (**Supplementary Fig. 3B**). Together, these capabilities enable robust morphological profiling of cellular heterogeneity across diverse staining panels (e.g. nucleus, ER, mitochondria, actin, and lysosomes) and cell types such as HEK293T cells and PBMCs (**Fig. 3D**).

**Fig. 3.**
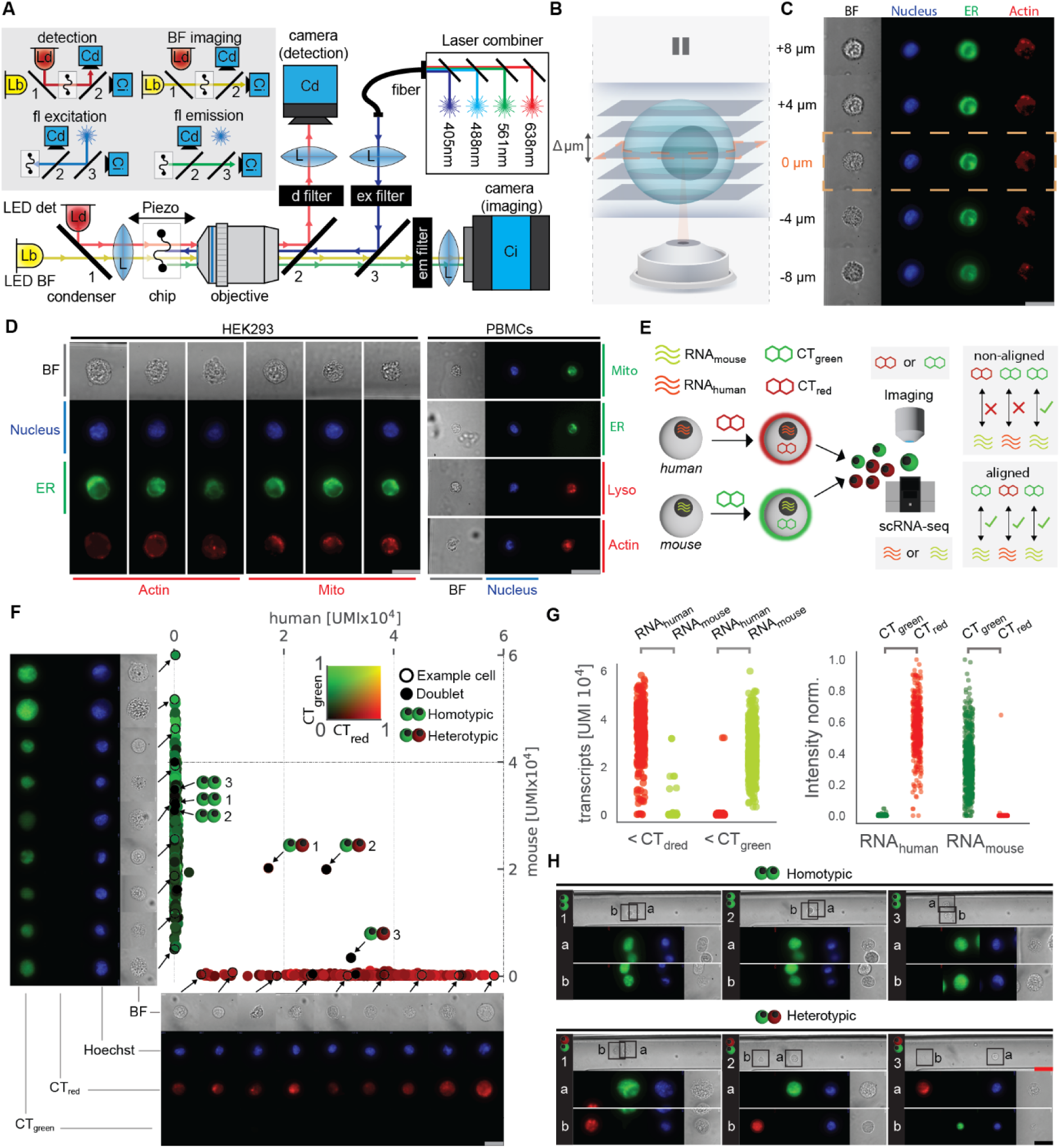
High-resolution single-cell imaging and image-transcriptome pairing in IRIS. **(A)** Simplified schematic of the microscope. The microscope is split into an imaging section and a detection section. For detection, a near-infrared LED (LED det, Ld) illuminates the chip and is imaged by the detection camera (Cd). For imaging, visible light illuminates the chip which is captured by the scientific-CMOS imaging camera (Ci). Both arms of the microscope are spectrally separated with shortpass dichroics (1, 2), and the detection light separated with a (d) filter. For fluorescence (fl) microscopy, the chip is illuminated with four laser lines, fiber-coupled to the microscope. The emission light is separated from the excitation light via a multiband dichroic (3), a multiband excitation (ex) filter, and a multiband emission (em) filter. The chip is coupled to a piezo actuator for focus adjustment. **(B)** Schematic of a single cell positioned in the stopping zone for high-resolution imaging with z-stack acquisition using a piezo-driven stage. **(C)** Representative z-stack images of a HEK293T cell in brightfield (BF) and three fluorescent channels: blue (Hoechst, nucleus), green (ER-Tracker, endoplasmic reticulum), and red (CellMask-Actin-DR, actin). Scalebar in grey corresponds to 20 μm. **(D)** Examples of morphological diversity in HEK293T cells (Hoechst, ER, mitochondria, actin) and PBMCs (Hoechst, ER, mitochondria, lysosomes, actin). Scalebars in grey correspond to 20 μm. **(E)** Species-mixing experiment to validate image to transcriptome alignment. Human HEK293T stained with CT-red and mouse NIH/3T3 stained with CT-green were processed on IRIS. The cell species was determined by sequencing and the identity of the stain by fluorescence microscopy. Pairing was evaluated by comparing both annotations. **(F)** Barnyard plot showing UMIs mapped to human vs. mouse genomes with points colored by CT-green/CT-red mean intensity measured inside image segmentation masks; representative single-cell images for each species are shown (arrow to empty circle). Using the imaging data of the encapsulation events that contained more than one cell (doublet events) were identified and highlighted (solid circle). Doublet events were annotated (homotypic, heterotypic) according to the fluorescence signal of each cell as shown in panel (H). Scalebar in grey corresponds to 20 μm. **(G)** Cross-modal correspondence between fluorescence intensity and transcriptomic species assignment at the single-cell level. *Left panel*: scatter plots showing the total number of UMIs (y-axis) mapped against each species’ genome stratified by fluorescence intensity categories, either higher fluorescence in the CT-red channel or in the CT-green channel (x-axis). For each intensity category, cells are split between *human* (left) or *mouse* (right) based on their transcriptomic profiles. *Right panel*: scatter plots showing fluorescence intensity (y-axis) in the green or red channel, stratified by transcriptomic species assignment (x-axis: *human* vs. *mouse*). **(H)** Manually selected, representative images of homotypic (numbered 1-3) and heterotypic (1-3) doublets identified by image analysis. Doublets were considered homotypic if they contained two cells with identical signal and heterotypic if they showed both red and green fluorescence. Scalebar in grey corresponds to 20 μm, in red to 40 μm.

To validate concordance between imaging and transcriptomics, we performed a human-mouse species-mixing experiment. HEK293T and NIH/3T3 cells were labelled with distinct live-cell dyes, CellTracker (CT)-red (*ex.* 638 nm) and CT-green (*ex.* 488nm), respectively (**Fig. 3E**). Barnyard plot showed clear species separation with minimal cross-contamination (**Fig. 3F**) and fluorescence intensities largely matched UMI-based transcriptomic species assignments: human cells exhibited high red channel intensity and human-specific transcriptomes, while mouse cells showed the converse (**Fig. 3G**). Morphological data also revealed that NIH/3T3 cells were slightly larger than HEK293T cells on average (**Supplementary Fig. 3E**), and allowed the identification of doublets (**Fig. 3H**). Transcriptionally defined mixed-species doublets were confirmed visually by co-localized green and red fluorescence, while homotypic doublets exhibited strong fluorescence in only one channel.

Together, these results demonstrate that IRIS achieves one-to-one matching of single-cell images with transcriptomes, enabling orthogonal validation of cell identity and direct measurement of morphological features such as size, subcellular staining patterns, and doublet status.

### Integrated imaging and transcriptomics reveal molecular correlates of continuous cell-cycle dynamics

As a first application, we profiled the cell cycle in Fluorescence Ubiquitin Cell Cycle Indicator^33,34^ (FUCCI)-expressing 3T3 fibroblasts. The cell cycle is continuous, well-characterized, and supported by a robust live-cell reporter (FUCCI), making it an ideal benchmark for validating the phenomics capacity of our IRIS platform. We profiled 5,191 cells, capturing FUCCI red (AzaleaB5-hCdt1, G1 phase) and green (h2-3-hGem, G2/M phase) as well as nuclear morphology (Hoechst 34580) (**Fig. 4A-B**).

**Fig. 4.**
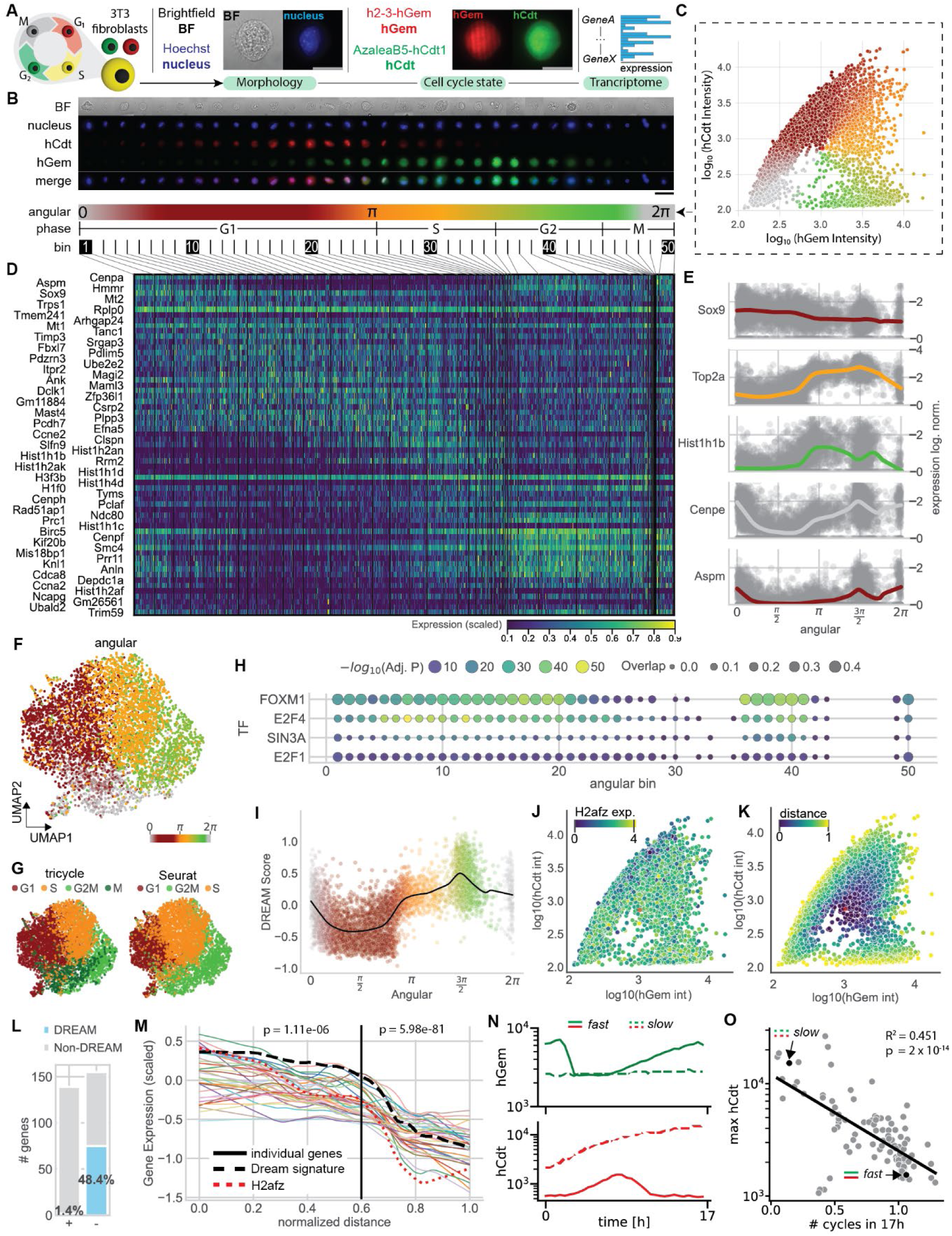
Phenomics analysis of the cell cycle in 3T3 fibroblasts. **(A)** Schematic of the experimental setup. A total of 5,191 3T3 cells expressing the FUCCI reporter were processed on IRIS. For each cell: (i) cell cycle state was assessed via the FUCCI reporter, (ii) nuclear morphology was captured by fluorescence imaging (Hoechst 34580), (iii) whole-cell morphology and shape were obtained by brightfield (BF) imaging, and (iv) transcriptomic profiles were measured. Scalebars in grey correspond to 20 μm. **(B)** Example cell images, ordered according to the angular score derived in (C). Scalebar in black in the lower right corner corresponds to 40 μm. **(C)** A continuous “angular score” was computed from FUCCI intensities, and cells were binned into 50 discrete angular intervals (shown on the left) and in Supplementary Fig. 4A. The color code is depicted on the left side of the figure (dashed arrow). **(D)** Expression of top three bin-specific genes per cell for differentially expressed (DE) genes calculated from the angular bins in (C). DE genes were computed for each individual angular bin (1-vs-all comparison, bins depicted above). Genes identified in multiple bins are only displayed once in the bin with the lowest index. **(E)** Per cell log-normalized gene expression of a manually selected subset of genes with varying expression dynamics as a function of the angular score (C). **(F)** UMAP embedding of all cells based on gene expression data, color-coded by angular score.**(G)** UMAP from (F), color-coded by Tricycle (left) and Seurat (right) cell cycle annotations. **(H)** Bubble plot showing transcription factor (TF) target gene enrichment based on DE genes from the angular bins. The top 4 cell cycle-related TFs are shown (a comprehensive overview is available in Supplementary Fig. 4E). Color code indicates Fisher’s exact test statistical significance, measured as -log10 adjusted p-value (Benjamini-Hochberg correction), dot size: the fraction of overlap between input list and TF target list. **(I)** DREAM complex target gene activity per cell as a function of the angular score, color-coded by angular color code as shown above. Solid line represents the DREAM complex target gene activity trend along the angular score (smoothed average).**(J)** FUCCI intensities as shown in (C) color-coded by *H2afz* log-normalized expression. **(K)** FUCCI intensities as shown in (C) color-coded by intensity distance metric (**Methods**). **(L)** Bar chart of DE genes with respect to the intensity distance metric in angular bins 18–22. Bars are colored by DREAM target gene representation. DE genes positively and negatively associated with the intensity distance metric are indicated with ‘+’ and ‘-’, respectively. **(M)** Line plots showing the expression of one cluster (cluster 9, Supplementary Fig. 4G) of downregulated DE genes along the intensity distance metric in angular bins 18–22. DREAM complex activity is overlaid as a dashed black line. **(N)** Fluorescence intensities of h2-3-hGem (green), AzaleaB5-hCdt (red) of two individual cells (one slow-cycling and one fast-cycling) that were tracked for 17h. The dashed line indicates the slow-cycling cell and the solid line the fast-cycling cell. **(O)** Maximum AzaleaB5-hCdt fluorescence intensity in relation to absolute angular difference per cell in 100 FUCCI 3T3 fibroblasts tracked over 17 h. A linear regression was fitted to the data. Two representative cells (slow, fast) shown in (N) are highlighted.

We first asked whether phenotypic and gene expression measurements could reciprocally inform cell cycle annotation and gene discovery. Unlike traditional methods such as flow cytometry-based sorting, IRIS profiles each cell at both the phenotypic and transcriptomic level, enabling a continuous read-out for every cell along the cell cycle (**Fig. 4C**). Based on the FUCCI intensities, we computed the cell cycle state of each cell by calculating a continuous ‘cell cycle angle’ score^35^ (**Fig. 4C**, see **Methods**) and grouped cells into 50 angular cell cycle bins (**Supplementary Fig. 4A**). We used this approach to capture gradual transcriptomic changes across the cell cycle and to avoid information loss because of coarse phase labels^36^. For each bin, we identified differentially expressed (DE) genes among the top 2000 highly variable genes (HVGs) (**Fig. 4D**, **Supplementary Fig. 4B**, **Methods**) with varying expression dynamics along the cell cycle (**Fig. 4E**). This yielded in total 670 cycling genes (33% of HVGs), of which 20% were part of Gene Ontology (GO) term “cell cycle”, including 46% of the 97 genes in the cell cycle reference panel curated by Tirosh et al.^37^, a set commonly used for annotating cell cycle phases in scRNA-seq datasets^38^ (**Supplementary Fig. 4C**). Interestingly, a substantial fraction comprising 527 genes was found in neither signature but was associated with Connective Tissue and Fibroblast tissue (**Supplementary Fig. 4D**), indicating that these are cycling genes that are cell type specific. Conversely, we compared our continuous angular annotation (**Fig. 4F**) to two transcriptome-only reference methods: Seurat’s 97-gene cell cycle panel based on Tirosh et al. and Tricycle^37,39^ (**Fig. 4G**), which projects scRNA-seq data onto a pre-trained continuous cell cycle trajectory to assign each cell a phase angle without any phenotypic information. Across all three approaches, cell cycle positioning was broadly concordant (**Fig. 4 F-G**). However, Seurat assigned a higher proportion of cells as S-phase compared to both Tricycle and our FUCCI-based angular annotation, consistent with prior observations^40^.

To demonstrate how IRIS can uncover new regulatory associations, we focused on transcription factors (TFs) whose target gene activity varied across cell cycle bins. We recovered known cell cycle regulators with phase-specific activity such as FOXM1^41^ (active during G1/S and G2/M transitions) and E2F4 (peaking in G1), as well as factors showing broader, continuous regulation including E2F1 and SIN3A (**Fig. 4H**, full set **Supplementary Fig. 4E**). Because E2F4 is a core component of the DREAM complex that represses G1/S transition genes during quiescence (G0), we next inferred DREAM complex activity using a published target gene signature^42^. DREAM complex activity peaked in late G1, reflected by down-regulation of its target genes (**Fig. 4I**). Bins with high DREAM activity were enriched for cell cycle genes (*Ccna2*, *Cdk1*) and chromatin regulators such as *H2afz*. Notably, *H2afz*, previously reported to be downregulated upon quiescence entry^43^, also showed lower expression in cells with high FUCCI reporter intensities (**Fig. 4J**). These observations suggest a functional link between FUCCI signal strength and the transition into G0, indicating that FUCCI intensity may capture a cell’s propensity to enter quiescence, beyond simply marking its position within the cell cycle.

To quantify this relationship, we defined a FUCCI ‘intensity distance’ metric along the cell cycle angular axis and correlated gene expression with this metric in each angular bin (**Methods**, **Fig. 4K**). Genes most strongly associated with FUCCI intensity clustered in the G1/S transition (window 21; **Supplementary Fig. 4F-G**). Notably, DREAM complex target genes were markedly enriched among negatively correlated genes (48% versus only 1.4% positively associated; **Fig. 4L**), reinforcing the link between high FUCCI reporter intensity and DREAM complex–mediated repression during late G1 and the G1/S transition. When examining the relationship between our intensity proxy and these anticorrelated genes (**Fig. 4M**), we observed a non-linear pattern with two statistically significant distinct domains of DREAM repression (details in **Methods**): an initial phase showing a significant, but weak negative association (norm. dist. < 0.6, p = 1.11e-06), followed by a sharp transition into a stronger negative association (i.e. stronger repression) (norm. dist. > 0.6, p = 5.98e-81). Quiescence-associated genes, such as *H2afz*, followed this biphasic pattern, suggesting that the stronger repressive regime corresponds to cells fully exiting the cycle into G0. In contrast, cells with a weaker association may correspond to cells in a “primed” state, i.e. slowing progression through late G1 without fully committing to quiescence.

To further examine how FUCCI reporter intensity reflects cell cycle dynamics, we performed live-cell imaging to monitor cell cycle progression in individual cells over time. If FUCCI reporter intensity reflects cell cycle dynamics, cells with higher AzaleaB5-hCdt1 (G1/red) signal should progress more slowly through the cell cycle or, in extreme cases, remain quiescent. We therefore tracked 100 individual 3T3 FUCCI cells over 17 hours (two exemplary tracks shown in **Fig. 4N**) and determined their cell cycle angle as above for each timepoint. From the absolute change in angle, we estimated each cell’s progression through the cell cycle over the 17-hour observation period (**Methods**), finding that most cells completed roughly one full cycle (**Fig. 4O**). Notably, cells that covered a smaller angular distance and thus progressed more slowly exhibited higher maximal AzaleaB5-hCdt1 intensity, consistent with prolonged G1 residence, as confirmed by linear regression (**Fig. 4O**). Taken together, these results suggest that DREAM complex target genes are downregulated in more slowly cycling cells. This implies that DREAM complex activity may influence not only quiescence regulation but also broader aspects of cell cycle dynamics.

These results demonstrate how IRIS uniquely bridges dynamic fluorescent reporters with genome-wide molecular profiling, enabling continuous cell cycle reconstruction without prior gating. By directly linking FUCCI reporter intensity to a cell’s transcriptional state at scale, IRIS revealed DREAM complex repression and cell cycle speed relationships that would likely remain obscured in single-modality analyses.

### Machine learning couples imaging and transcriptomics across single cells

To further leverage IRIS’s capacity of pairing cellular morphology with matched transcriptomes at a large-scale, we tested whether this paired data can be used to uncover correlations between morphology and gene expression. We laid our main emphasis on retaining biological interpretability and for this reason only employed comparably simple machine learning models with single modality inputs. We trained a ResNet18^44^ convolutional neural network (CNN) with regression outputs to predict expression of the top 2,000 HVGs from two inputs: nuclear morphology (Hoechst) and whole-cell shape masks from brightfield images (**Fig. 5A**). We trained the models to minimize the mean squared error (**Methods**), yielding the conditional expectation of gene expression given the input images. We evaluated performance on a set of test cells that were not observed during training (20% of total cells, **Methods**). The nuclear model probed information-rich organelle structure, and its input images were min–max normalized so predictions could not rely solely on raw intensity values. In contrast, the mask model prioritized interpretability by operating on binary whole-cell masks, from which intuitive features such as size and circularity are easy to derive. As a benchmark, we built a reference model predicting gene expression from the continuous cell cycle angular score, providing both a performance ceiling and biological ground truth.

**Fig. 5.**
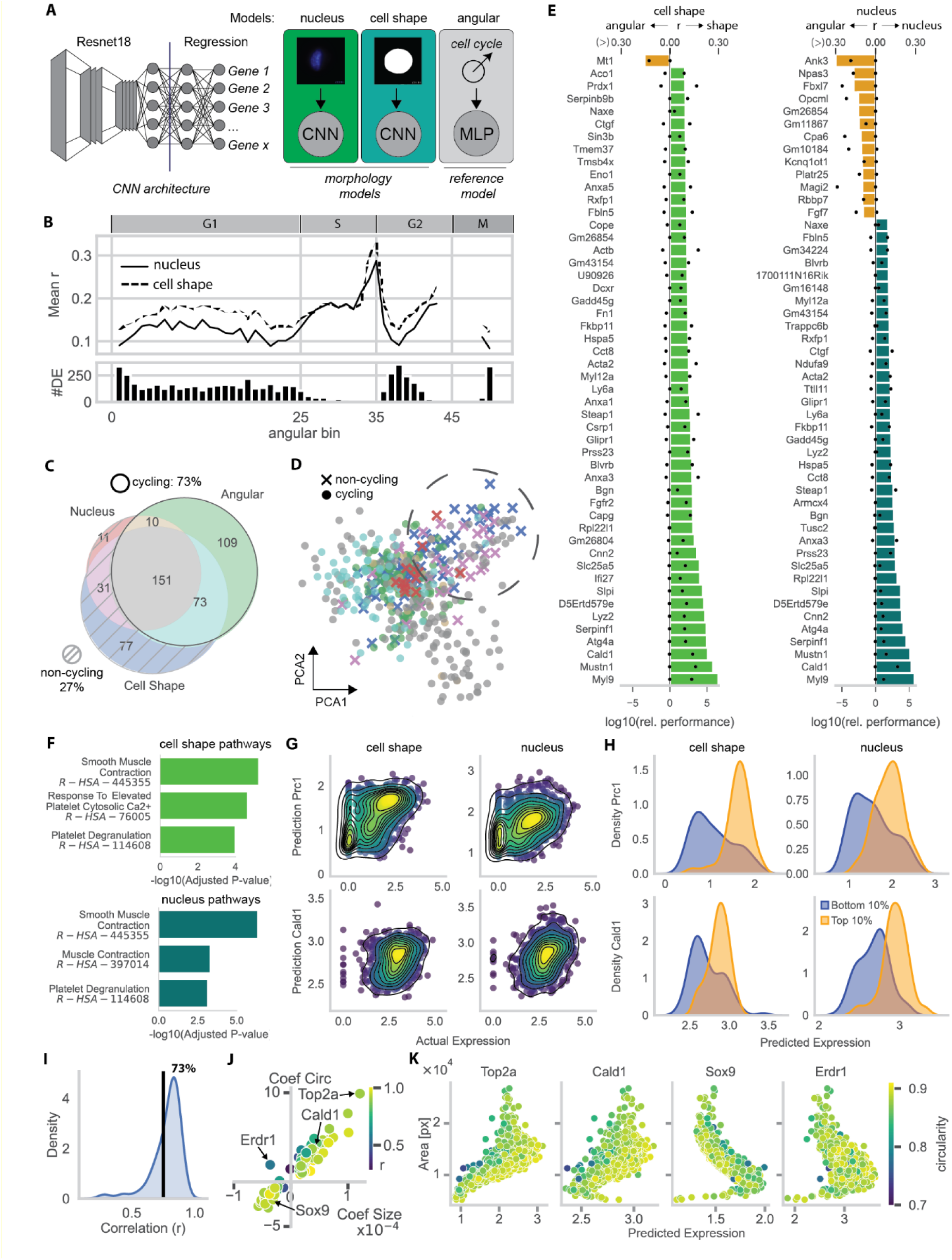
Machine learning models for predicting gene expression from cell morphology and angular features. **(A)** Conceptual overview of the three models that were trained. Two image-based convolutional neural networks (CNNs, ResNet18 backbone with regression layer) were trained using either cell segmentation masks (cell shapes) or Hoechst-stained nuclear images as input to predict gene expression. A third model, a multilayer perceptron (MLP), was trained to predict gene expression from the angular score. **(B)** Model performance for cell cycle associated genes. *Top*: mean Pearson correlation of predicted versus observed gene expression values across differentially expressed (DE) genes of angular score bins (as in Fig. 4). *Bottom*: number of DE genes contained in each bin. **(C)** Venn diagram of predictable genes (adjusted p-value < 0.01, Pearson p-value corrected with Benjamini–Hochberg method). All genes predictable by the angular model were classified as ‘cycling’. **(D)** Principal component analysis (PCA) of all genes that were predictable with statistical significance (adjusted p-value < 0.01) by either model (angular, cell shape, nucleus). Genes are colored according to the Venn diagram categories defined in panel (C). Note: each dot represents one gene. **(E)** Prediction performance of the cell shape and nuclear models relative to the angular model. Bar plots show relative performance; dots indicate absolute Pearson correlations (angular model values shown on the left, respective models on the right). Top 50 genes with highest or lowest relative performance in each model are shown. **(F)** Gene set enrichment analysis (GSEA) of genes that had 2-fold or higher relative predictive performance int the cell shape model or the nuclear model, respectively. **(G)** Scatter plots of predicted versus observed expression for two genes that were among the most predictable cell shape and nuclear model’s genes. *Prc1* was selected as a representative for cycling genes, *Cald1* was selected as a myofibroblast marker. Colored by point density. **(H)** Density distributions of predicted expression values for the top and bottom 10% of cells ranked by observed (actual) expression. **(I)** Density distribution of correlation values between predicted expression and morphological features for the genes shown in panel (J). **(J)** Regression coefficients of a linear regression model using cell size and circularity to explain predicted expression values from the cell shape model. The x-axis depicts the Size coefficient, the y axis the Circularity coefficient. Results are shown for the top 100 predictable genes of the cell shape model. The color-code represents the Pearson correlation coefficient. **(K)** Scatter plot of predicted expression values from the cell shape model against cell size (area), color-coded by circularity.

For cell cycle-related genes, the angular reference model identified 343 significant (Pearson p-value < 0.01, FDR corrected) associations, peaking at a test Pearson correlation of 0.76 for the canonical cycling cell marker *Top2a*. The nuclear and cell shape models captured 203 and 332 associations, respectively, with maximum correlations of 0.37 (*Top2a*) and 0.38 (*Prc1*, a cell cycle regulator) (top 100 genes in **Supplementary Fig. 5A**). Mean performance for the top 100 best predicted genes by Pearson correlation of the morphology models, i.e. cell shape and nucleus, reached 39% and 47% of the reference, respectively, but phase-specific analysis revealed distinct strengths. Specifically, we used the angular bin-specific DE genes from **Fig. 4D** as phase-defining gene sets, and for each angular bin calculated the mean Pearson correlation of actual and predicted expression (**Fig. 5B**). This revealed that the cell shape model performed better than the nuclear model during early-to-mid G1 (bins 0-25), likely reflecting growth-associated size changes, while the nuclear model displayed increasing average associations in the S phase, consistent with the structural reorganization accompanying DNA replication.

We next asked whether the morphology models capture additional biological processes beyond the cell cycle. Comparing all significantly associated genes (adjusted p-value < 0.01) across all three models revealed that 74% were cell cycle-related, of which 68% were shared with the angular model, while the remaining 26% were uniquely discovered by the morphology models (**Fig. 5C**). This suggests that their association is independent of cell-cycle influences. To determine whether the remaining genes were biologically distinct from cycling genes, we performed dimensionality reduction on significantly associated genes (**Fig. 5D**). This analysis showed that non-cycling genes formed a discrete cluster, distinct from most cycling-associated genes. Aiming to find a common biological function, we ranked all significantly associated genes of both morphological models relative to the angular model (**Fig. 5E**), and performed GSEA on genes with > 2-fold-better relative performance. Both models (nuclear and mask) converged on similar enrichments, notably muscle contraction pathways (**Fig. 5F**). In the fibroblast context, this points to the presence of an activated myofibroblast subpopulation in our cell cultures, supported by high expression of smooth muscle actin (*Acta2*) and strong associations with contractility-related genes such as Caldesmon1 (*Cald1)*. To better understand how the models predicted distinct biological states, we examined the top cell cycle gene *Prc1* and the myofibroblast-linked gene with the strongest association, *Cald1* (**Fig. 5G**). For both, predicted expression tracked measured values linearly with low batch effects (**Supplementary Fig. 5B**). As expected, the distribution of predictions was narrower than the actual expression distribution, because the conditional expectation of gene expression pulls extreme observations toward the average reducing the spread relative to the observed data. We then tested whether the morphology models could reliably separate high- and low-expressing cells for these marker genes, revealing that both models showed robust discrimination (**Fig. 5H**). Taken together, these findings suggest that supervised machine learning can uncover biological processes directly from paired cell morphology and gene expression data.

To gain interpretability, a long-standing challenge with black-box neural networks, we focused on two ‘simple’, easy-to-compute shape descriptors: cell size, a hallmark of growth, and cell circularity (**Methods**). We fit linear models to the CNN-based predictions for the top 100 cell shape-associated genes and found that most were relatively well-explained by the two descriptors with 73% of all gene predictions by the linear regression models reaching a Pearson r higher than 0.75 (**Fig. 5I**). With the exception of *Erdr1*, genes fell into two categories: positively correlated with size and circularity such as *Top2a*, or negatively correlated with both, such as *Sox9* (**Fig. 5J**). Inspection of both gene groups, including *Cald1*, confirmed non-linear associations with CNN-based predictions (**Fig. 5K**). Strikingly, *Top2a* and *Cald1*, while having distinct biological functions, were predicted from the same morphological feature with similar dynamics, revealing that shared phenotypic descriptors can encode different transcriptional programs. These analyses highlight a capability that is non-trivial: IRIS can resolve subtle, non-discrete cell states and uncover hidden biological processes by directly coupling image-derived morphology with high-resolution transcriptomes.

### Morphological architecture reveals distinct transcriptional programs in naïve CD8⁺ T cells

So far, we have used IRIS to uncover distinct transcriptional programs and link them to corresponding morphological phenotypes. Next, we aimed to determine whether IRIS can resolve transcriptomic differences that arise between distinct morphological forms within the same cell type. Specifically, we set out to assess whether structured cell features reflect functional molecular signatures. We focused on peripheral blood mononuclear cells (PBMCs), a morphologically heterogeneous population^45^ in which both lymphoid and myeloid subsets display well-documented differences in nuclear architecture, including differences in shape, lobulation and the presence of deep invaginations^46,47^. This made PBMCs an ideal system to investigate whether variation in nuclear shape alone could delineate transcriptionally distinct states.

Using IRIS to pair nuclear morphology with transcriptomic profiling in PBMCs (n=3,033) from four healthy donors, we recovered all major lymphoid and myeloid subsets based on transcriptomic identity (**Fig. 6A**), validating the platform’s ability to resolve molecularly distinct immune cell populations with high fidelity from mixed samples. Having established this baseline, we next asked whether IRIS could uncover additional morphological heterogeneity within transcriptionally defined subsets. Specifically, we stratified cells by the presence of nuclear invaginations, reasoning that these structural variations in nuclear morphology might mark further molecular diversity not captured by standard subset definitions. This revealed characteristic invaginated nuclear morphologies across both myeloid and lymphoid subsets (**Fig. 6B-C**).

**Fig. 6.**
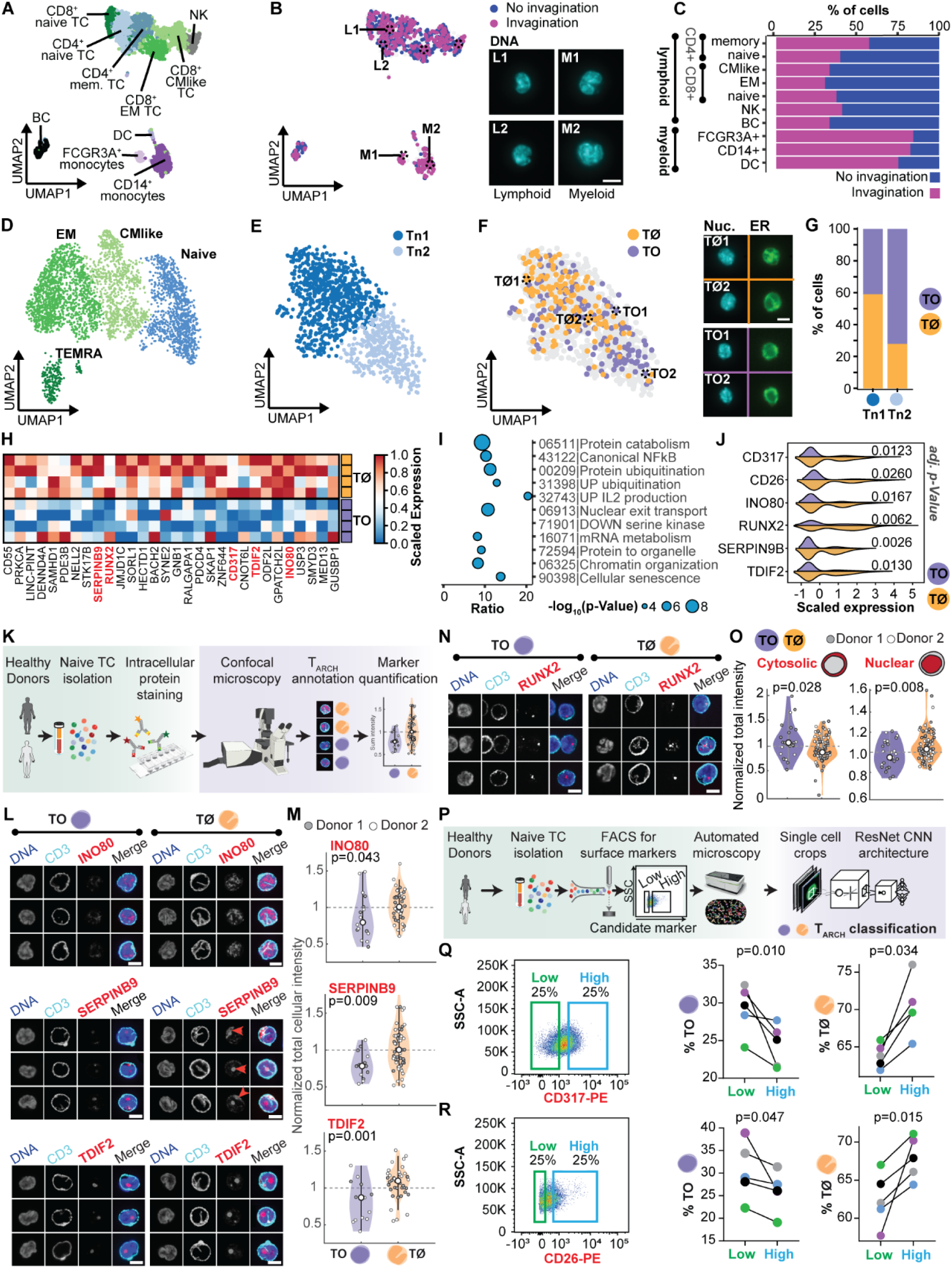
T cell architecture profiling with IRIS links nuclear-ER organization to activation-associated transcriptional programs in naive T cells. **(A-C)** IRIS profiling of PBMCs combining nuclear morphology imaging with scRNA-seq. **(A)** UMAP of integrated single cell PBMC transcriptomes (n=3,033) from four donors. **(B)** Nuclear morphology classification via manual curation of invaginated vs. non-invaginated cells, overlaid onto the PBMC scRNA-seq UMAP: invaginated (n=289) vs. round (n=380) nuclei with representative images (n=2,364 ambiguous). Scale bars, 5 μm. **(C)** Proportions of nuclear morphologies within each PBMC subset. **(D-J)** Peripheral blood-derived CD8^+^ T cells (n = 3,969) from four donors were profiled with IRIS, pairing ER and nuclear morphology with scRNA-seq. **(D)** UMAP of integrated CD8^+^ T cells from four donors. Cells are colored based on the different T cell subsets including naïve, CMlike, TEMRA and EM subset. CMlike, central-memory-like T cells | TEMRA, CD45RA^+^ effector-memory | EM, CD45RA^-^ effector memory. Canonical markers for subsets in Supplementary Fig. 6A. **(E)** Subset-specific UMAP re-embedding of naïve CD8⁺ T cells (Tn1, naïve CD8+ T cell subset 1, Tn2, naïve CD8+ T cell subset 2). **(F)** TØ (‘stripy’, ER displays a stripe-like pattern) and TO (‘conventional’, ER forms a continuous layer surrounding the nucleus) architectures in T cells, superimposed on re-embedding of only the naïve CD8^+^ T cells (Tn). *Right*: representative images of each morphology linked to their UMAP position. Scale bars, 5 μm. **(G)** Proportions of TØ and TO cells within the two naïve T cell subsets. **(H)** Top 30 differentially expressed genes between TØ or TO naive CD8^+^ T cells, shown as mean-centered mini-bulk expression per donor. Significance determined by Wilcoxon test, Benjamini-Hochberg correction applied. Genes validated at the protein level are indicated in red (panel K-R). **(I)** GO terms enriched among genes upregulated in TØ versus TO naive CD8+ T cells. **(J)** Expression of selected upregulated genes in TØ versus TO naive CD8+ T cells across all naive T cells. Significance determined by Wilcoxon test, Benjamini-Hochberg correction applied. **(K)** Workflow for protein-level validation by confocal microscopy. **(L)** Representative INO80, SERPINB9, and TDIF2 stainings in TØ and TO cells from two donors; single z-planes are shown for CD3 marking the nuclear invagination, Hoechst (DNA) and the respective protein. ‘Merge’ shows average CD3^+^ and protein abundance. Scale bars, 5 μm. **(M)** Quantification of INO80, SERPINB9, and TDIF2 protein abundance in TØ versus TO cells across the two donors (two-sided t-test). **(N)** Representative RUNX2 stainings in TØ and TO cells. Single z-planes are shown for CD3 marking the nuclear invagination, Hoechst (DNA). ‘Merge’ shows average CD3^+^ and protein abundance. Scale bars, 5 μm. **(O)** Quantification of cytosolic and nuclear RUNX2 protein abundance in TØ and TO cells (two-sided t-test) across the two donors. **(P)** Workflow for surface marker-based enrichment of TØ cells. **(Q-R)** FACS gating for low/high expression of indicated markers and corresponding enrichment of TØ cells. **(Q)** CD317; **(R)** CD26. Scatterplots show baseline-normalized proportions of TØ among sorted from cell originating from five individual donors (two-sided t-test).

Invaginated nuclei in T cells have previously been linked to T cell antigen response strength, and subsequent differentiation^46^. To examine how transcriptional states relate to this morphological architecture, we focused on molecularly resolving two T cell morphological classes within naïve CD8⁺ T cells: (1) TØ ‘Stripy’ cells, characterized by ER enrichment within deep nuclear invaginations, and (2) TO ‘Conventional’ cells, in which the ER is uniformly distributed around smooth, non-invaginated nuclei^46^. Of note, single-cell resolved transcriptional profiling of these distinct morphologies has so far been unattainable due to methodological limitations in image resolution and the lack of scalable cell selection strategies, highlighting the core technological gap that IRIS was designed to overcome. In a dedicated experiment profiling 3,969 peripheral blood-derived CD8+ T cells from four healthy donors (**Methods**), we applied IRIS to capture both nuclear+ER morphology and transcriptome at single-cell resolution. The major CD8⁺ T cell subsets, including naïve, effector-memory (EM), central-memory-like (CM-like), and CD45RA^+^ effector-memory (TEMRA) T cells were readily identifiable based on their transcriptional profiles (**Fig. 6D**, **Supplementary Fig. 6A-B**) with their proportions varying across healthy donors (**Supplementary Fig. 6C**). Notably, among naïve CD8⁺ T cells, TØ cells were enriched in a specific subset (Tn1; **Fig. 6E-G**). However, the association was not absolute, making naïve CD8⁺ T cell subset-based differential expression inherently noisy. To resolve the molecular programs underlying these distinct morphologies, we leveraged IRIS’s ability to directly stratify individual cells by their morphology, allowing differential expression analysis between TØ and TO cells. This revealed higher expression of TCR signalling modulators (e.g. RUNX2, SKAP1), chromatin remodelers (e.g. INO80, TDIF2), transcriptional regulators (e.g., LINC-PINT), surface markers (e.g., CD55, CD317), and organelle organization genes (e.g. SERPINB9, SYNE2) (**Fig. 6H**, **Supplementary Fig. 6D**). The upregulation of genes linked to an activated state of the profiled naïve T cells was consistent with prior observations that TØ cells respond rapidly to TCR stimulation^46^, further supported by enriched gene ontology terms such as mRNA metabolism, IL-2 production, and canonical NFκB regulation (**Fig. 6I**).

The discovery of a distinct transcriptional program associated with TØ architecture prompted us to ask whether these molecular differences are also evident at the protein level. To address the differential protein expression associated with TØ morphology within the CD8⁺ naïve compartments, we performed immunofluorescence staining coupled to confocal microscopy for selected upregulated genes (**Methods**; **Fig. 6J-K**). We thereby focused our validation on genes whose functions are directly consistent with the distinct nuclear-ER architecture observed in TØ cells and with their observed activation propensity^46^. Specifically, we selected INO80 and TDIF2 (*DNTTIP2*), two chromatin remodelling proteins implicated in transcriptional accessibility and nuclear organization^48,49^, as well as SERPINB9, a protease inhibitor essential for early granule formation in cytotoxic T cells^50^ and RUNX2, a TF critical for T cell activation and tonic TCR signaling^51^, both of which were upregulated in the TØ subset and consistent with a heightened activation state. We found that protein levels of INO80, TDIF2, and SERPINB9 were higher in CD8⁺ naïve TØ cells, with nuclear enrichment of INO80 and TDIF2, and localization of SERPINB9 to the nuclear invaginations (**Fig. 6L-M**, **Supplementary Fig. 6E**). Furthermore, RUNX2 displayed differential nuclear localization, consistent with recent TCR activation (**Fig. 6N-O**), potentially linked to heightened tonic TCR signalling previously observed in TØ cells^52,46^.

Finally, we validated surface markers associated with TØ morphology within the CD8⁺ naïve compartment, identifying CD55, CD26 and CD317 as upregulated in TØ cells (**Fig. 6H**, **Fig. 6J, Supplementary Fig. 6D**). Because CD55 was broadly expressed across naïve CD8^+^ T cells and showed more limited discriminatory power (**Supplementary Fig. 6F**), we focused on CD317 and CD26 as more selective candidates (**Fig. 6H**, **Supplementary Fig. 6D**). We performed FACS enrichment followed by CNN-based morphological classification^46^ (**Fig. 6P-R**, **Supplementary Fig. 6G**, **Methods**) which revealed across five donors that naïve T cells with robust marker expression (top 25%) of CD317⁺ (encoded by *BST2*) or CD26⁺ (encoded by *DPP4*) exhibited significantly increased proportions of TØ cells (mean 6.5% ± 3.5 (p-value = 0.034) and 5.5% ± 4.0 (p-value = 0.015)), respectively, **Fig. 6Q-R**). These findings demonstrate that TØ identity is not only morphologically and transcriptionally defined, it is also cell surface-accessible, enabling prospective enrichment of architectural subsets.

By uniquely linking high-resolution imaging with deep molecular profiling, IRIS molecularly defined a previously reported but transcriptionally uncharacterized axis of T cell heterogeneity shaped by nuclear and ER architecture. These results establish cellular morphology as a quantitative and functionally informative readout of immune cell state, demonstrating how integrated single cell phenomics can reveal hidden layers of immune regulation and connect structural variation to molecular function.

## Discussion

Cellular function emerges from the interplay of molecular, structural, and physical properties, reflecting a principle now recognized as the cellular dogma^53,54,5^. By enabling integrated molecular and image-based readouts at single-cell resolution and scale, IRIS directly addresses the need for technologies that can resolve these interdependent layers of cellular organization. Our proof-of-concept studies illustrate the value of this integration. In the cell cycle, IRIS recapitulated canonical dynamics while revealing regulatory associations with progression speed that emerged only through large-scale phenomic profiling. In T cells, phenotypic stratification by nuclear and ER morphology uncovered coordinated expression programs that were invisible to single-modality approaches^46^, and experimental validation supported these associations. These findings highlight how stratifying cells by phenotype can uncover otherwise difficult-to-access molecular programs, underscoring the interpretive power of integrated readouts.

Beyond descriptive phenotyping, the scale of IRIS datasets establishes a rich resource for systematically exploring how cellular morphology reflects molecular state. This represents an emerging frontier: while prior efforts relied on computationally linked datasets^55^ or coarse cell-type proxies^56^, IRIS directly provides ground-truth pairings at single-cell resolution. Thus, these datasets now make it possible to quantify how much phenotypic information captured across imaging modalities encodes transcriptional identity and regulatory activity. This is an inherently complex task, as morphology and gene expression represent complementary yet only partially overlapping information spaces. IRIS directly probes the intersection of these spaces, offering the data foundation needed to model how structure and state co-vary. This opens the possibility that, in well-characterized contexts, molecular profiles could eventually be inferred rather than directly measured, accelerating both discovery and translational applications.

To explore this potential, we examined how different imaging modalities capture molecularly relevant variation. We found that prediction fidelity depends strongly on the nature of the captured phenotype: FUCCI intensities yielded robust predictions but are inherently restricted to cell cycle dynamics. Hoechst and ER staining captured richer organelle-specific features, linking to broader molecular programs but also introducing interpretive complexity, since distinct transcriptional states can converge on similar morphological signatures. Brightfield imaging, by contrast, aggregates optical signals from all cellular components, providing a holistic but also highly convolved view of cell state. Rather than a limitation, these differences highlight how IRIS can provide benchmarks, allowing systematic comparison of which imaging modalities, spatial resolutions, and analytical frameworks most effectively capture molecularly meaningful variation. As imaging technologies continue to advance in spatial, spectral, and volumetric (3D) resolution^57^, we anticipate that IRIS will play a major role in unlocking the full potential of morphology-molecular inference.

IRIS is also inherently well suited for future explorations that probe cellular dynamics. Its continuous operation could, in principle, make it possible to monitor how perturbations reshape molecular and phenotypic states over time^21^, enabling cellular response applications in drug screening, immune monitoring, and dynamic cell biology. Moreover, because cells are transiently immobilized during imaging and processing, the platform’s modular design makes it conceptually compatible with a wide range of image modalities and downstream biochemical assays beyond scRNA-seq. This adaptability could eventually support new capabilities such as image-based sorting to prospectively isolate rare or functionally distinct morphotypes.

At present, several limitations remain: (i) throughput (∼1,000 cells per hour) must increase to reach population-scale studies. (ii) IRIS profiles cells in suspension, which removes spatial context and may alter features present in adherent or tissue environments. While this constraint is irrelevant for naturally circulating cells, it defines a complementary niche to spatial transcriptomics, focusing on deep molecular and morphological profiling at the single-cell level. (iii) Imaging resolution can also be further improved, as higher spatial detail and dimensionality will likely reveal deeper layers of heterogeneity and strengthen morphology–molecular inference. Finally (iv), as with any new hardware, widespread adoption will depend on engineering refinements and community access, though efforts are already underway to broaden availability.

Taken together, IRIS provides a versatile platform for dissecting the molecular and phenotypic dimensions of cell identity. Beyond serving as a profiling tool, it establishes a generalizable framework for translating between imaging and transcriptomics, thereby enabling direct exploration of the ‘cellular dogma’. This perspective reframes how cell biology can be studied, opening new avenues to systematically link morphology with molecular logic. While the current study focuses on proof-of-concept applications, the ability to build image-based predictors calibrated against transcriptomic ground truth suggests strong translational promise. This includes scalable blood-based assays where imaging is accessible and rapid but interpretability remains limited, setting the stage for advances in diagnostics, therapy monitoring, and precision medicine.

## Supporting information

Supplementary Table Antibodies

Supplementary Table Primers

---

**Supplementary Fig. 2.**
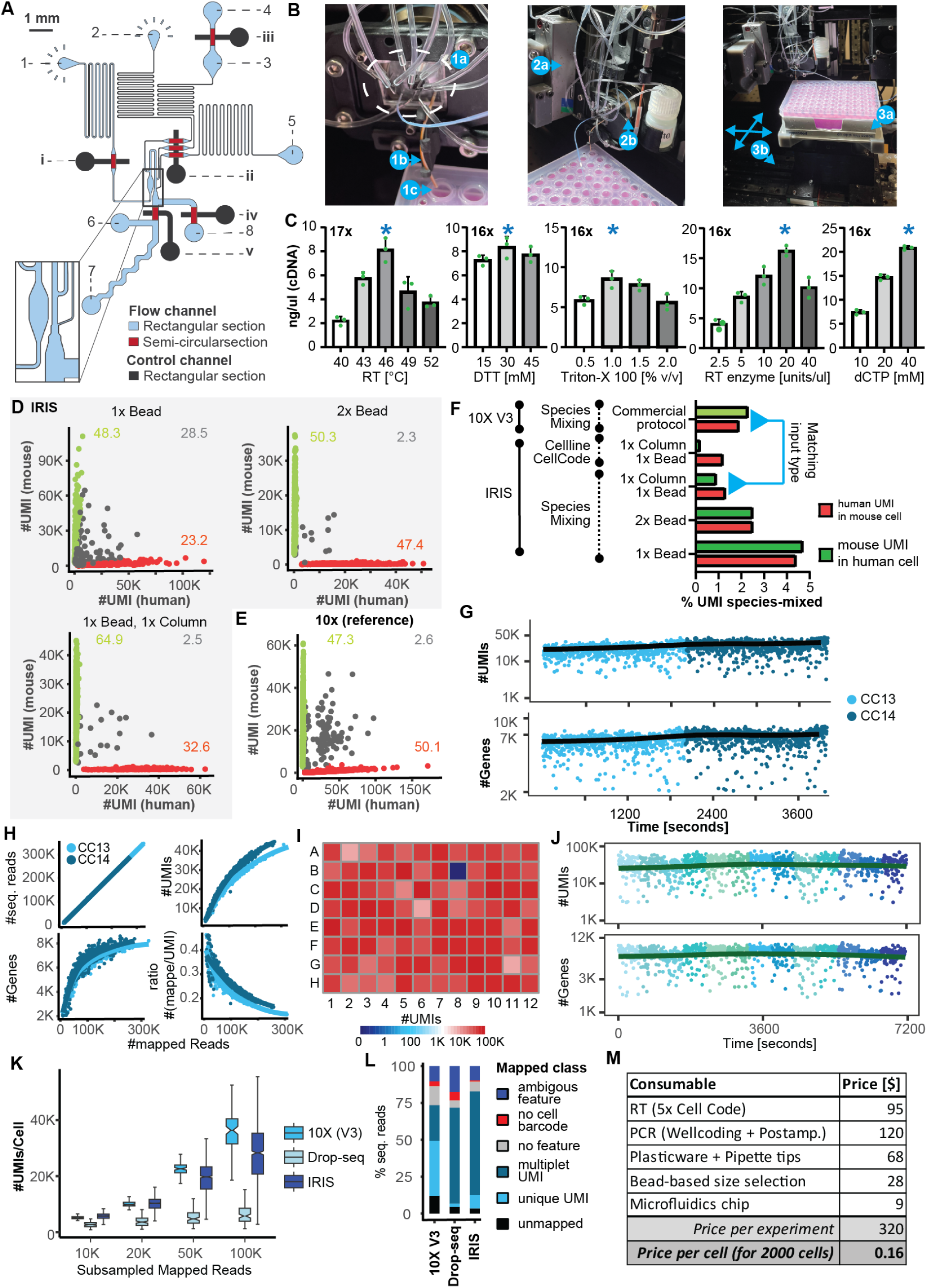
Optimization and validation of the deterministic ‘droplet consortia’-based workflow. **(A)** Dual-layer microfluidic chip with Quake-valve-controlled stopping, RT solution exchange, and encapsulation. Flow layer: semicircular (red) and rectangular (blue) cross-sections; control layer (black): push-up valves. Key inlets/outlets and valve functions are numbered as follows: 1) input channel for cell loading, 2) injection inlet for precise cell placement and stopping, 3) inlet for the “cell barcode” (CC)-containing reverse transcription (RT) solution, 4) outlet for the RT solution exchange, 5) oil inlet for droplet generation, 6) oil inlet for droplet ejection, 7) droplet mixing and exit channel, 8) liquid waste exit channel; i) cell-loading valve, ii) injection valve, which enables nanoliter-precision injection of liquid for cell placement, RT solution and oil injection for encapsulation, via a controlled over-pressurization of the valve (leak-flow), iii) RT solution exchange valve, iv) liquid waste outlet valve, v) droplet collection outlet valve. **(B)** *Left*: Picture of the microfluidic chip (1a) in the chip holder, all inlets and outlets connected with tubing, and the outlet capillary (1b) positioned over a well (1c). *Middle*: Picture of the same chip with the chip holder connected to the piezo actuator (2a), and the objective at the backside of the chip (2b). *Right*: Picture of the system including the plate mounted on a cool block (3a) which resides on the plate XY stage (3b). **(C)** Effect of RT temperature, DTT, Triton X-100, RT enzyme, and dCTP concentration on cDNA yield, as measured using one HEK293T cell per well, multiplexing 96 HEK293T cells. Stars indicate optimized conditions. Numbers indicate PCR cycles performed to amplify dual-indexed cDNA of 96 HEK293T cells. **(D)** Species-mixing (human HEK293T, red; mouse NIH/3T3, green) benchmark showing percentage of mixed-species UMIs after different well-coding primer cleanup strategies: 1x bead-based size selection (top left), 2x bead (top right), 1x column+1x bead (bottom left). Numbers indicate the percentage of total cells processed annotated with the respective species with the cutoff for mixed species cells (grey) being >10% mixed UMIs. **(E)** Mixed-species results from 10X Genomics 3’ V3 for comparison. Numbers and colors are as in (D). **(F)** Cross-species UMI percentage for cells with <10% mixed UMIs. Species mixing experiments (SM) were either performed with 10X Genomics 3’ V3 (10X V3) or with IRIS. Species-mixing design to assess droplet integrity ‘Cellline Cellcode’, where human HEK293T and mouse NIH/3T3 cells, each with a designated CC, were combined in the same well. **(G)** Continuous processing of HEK293T cells: number of UMIs (top) and genes (bottom) per cell over time, colored by CC RT primer (**Supplementary_Table_PRIMER**). **(H)** scRNA-seq quality metrics as mapped reads vs number of sequenced reads, genes per cell, detected UMIs per cell or ratio of UMI to mapped reads per cell, colored by CC RT primer. **(I)** Heatmap of number of UMIs across wells in a representative 96-well plate. **(J-L)** HEK293T cells were processed on IRIS for native multiplexing in ‘droplet consortia’ of 13 cells per well by introducing iteratively 13 different RT solutions containing different CCs. **(J)** UMI and gene counts per CC RT primer in native multiplexing of 13-cell ‘droplet consortia’. For the color code, see Fig. 2H. **(K)** UMIs per cell vs. downsampled mapped reads for IRIS, Drop-seq (on DisCo) and 10x Genomics 3’ V3. **(L)** Proportion of sequenced reads passing demultiplexing and sequencing quality specifications to all mapping categories from HEK293T cell scRNA-seq across indicated methods. **(M)** Details the estimated run cost of one IRIS experiment with 2000 cells across the whole workflow.

**Supplementary Fig. 3.**
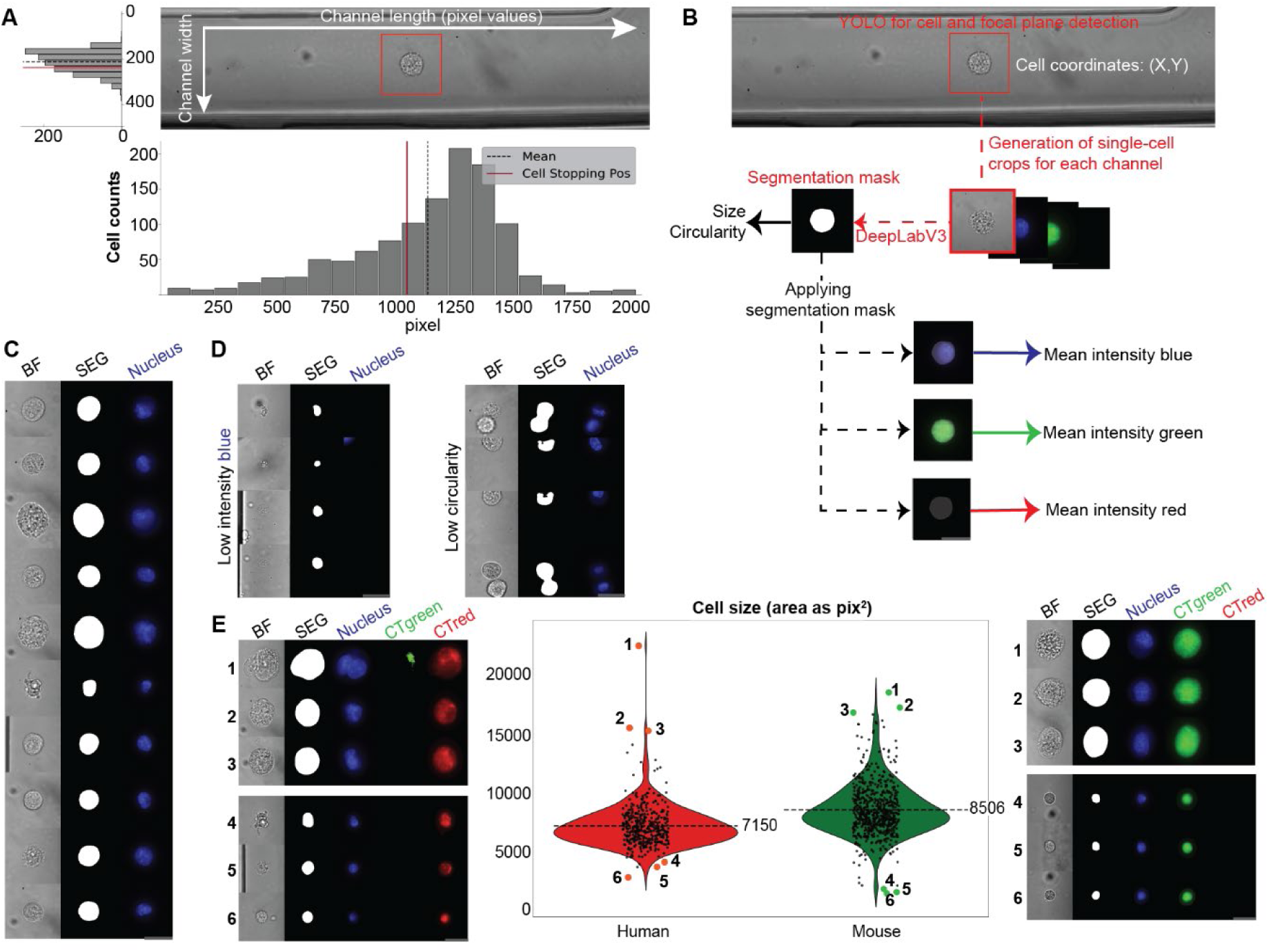
Characterization and quality control of the IRIS single-cell imaging pipeline. **(A)** Distribution of stopping positions along the channel length (bottom) and width (left), within the imaging zone in a representative species-mixing run (corresponding to Fig. 3F); dashed line indicates the mean stopping position, red mark highlights the example cell. **(B)** Pre-processing workflow. Cell coordinates are determined with YOLO and are used to generate centered image crops (250×250 pixels). DeepLabv3 segmentation masks are applied to extract cell size, circularity, and mean fluorescence per channel, while excluding background signal. **(C)** Examples of BF images (left), segmentation masks (middle), and Hoechst channel (right) illustrating accurate cell boundary detection across diverse morphological conditions. **(D)** Representative outliers identified during pre-processing: cells lacking Hoechst signal (left) or with low circularity (<0.5), often corresponding to debris, doublets or partial cell captures (i.e., cut off at the edge of the stopping zone in the microfluidic channel) (right). **(E)** Violin plots of segmented cell areas by transcriptomic species identity (corresponding to Fig. 3F). Example images of three small and three large cells with HEK293T cells (left) and NIH/3T3 cells (right). Cells lacking Hoechst signal or with low circularity (<0.5) were excluded to ensure data quality. Scalebars in grey correspond to 20 μm.

**Supplementary Fig. 4.**
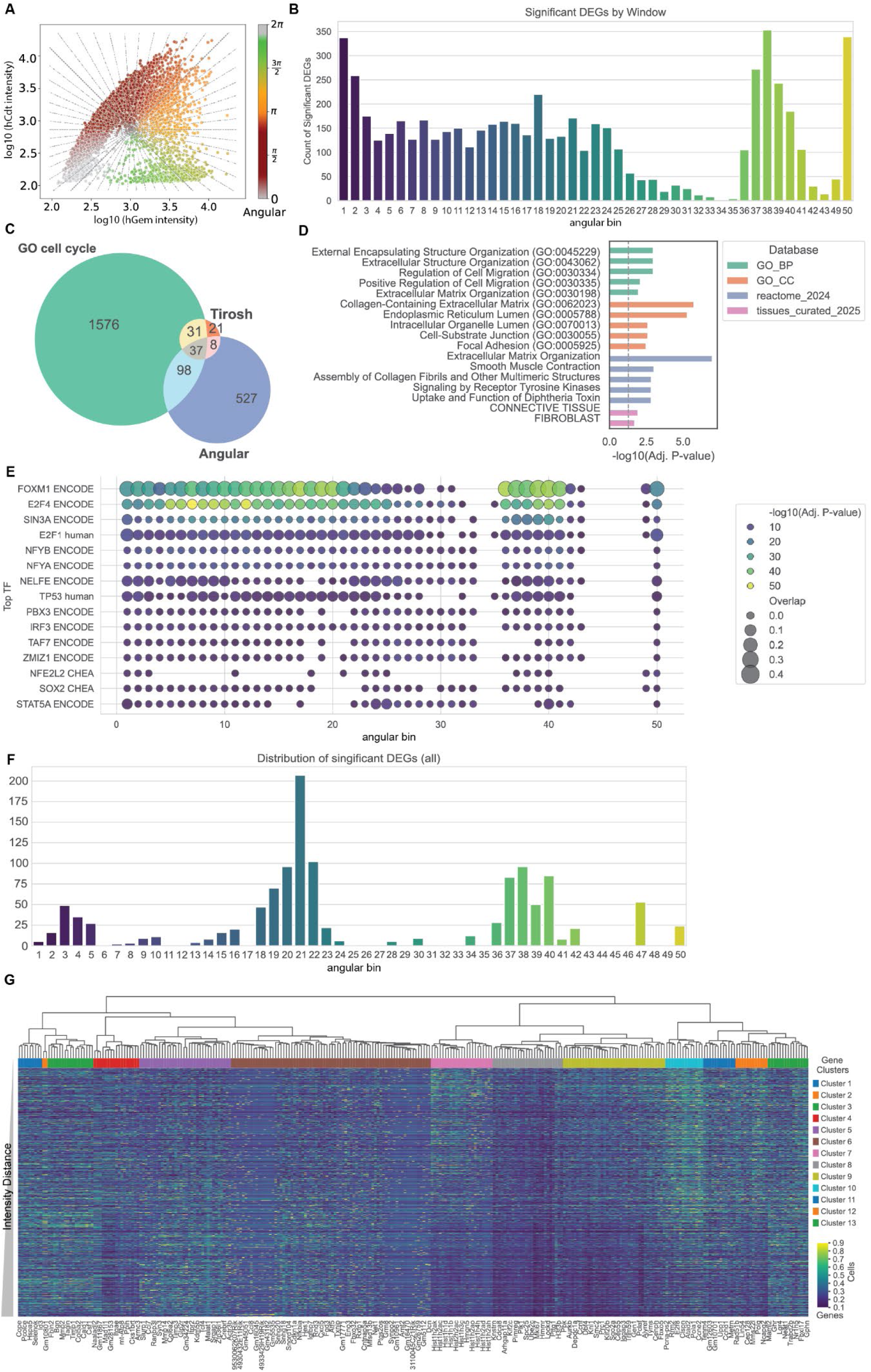
Phenomics analysis of the cell cycle in 3T3 fibroblasts. **(A)** Intensities as shown in Fig. 4C overlaid with angular bin lines. **(B)** Number of differentially expressed (DE) genes identified per angular bin/window. **(C)** Overlap of identified cell cycle-associated genes by the angular score (*Angular*) and cell cycle signature genes (*Tirosh* and *GO cell cycle*). **(D)** GSEA results of genes considered non-cycling (not in signatures shown in C). **(E)** Transcription factor target gene enrichment based on angular bin DE-genes. **(F)** Number of FUCCI intensity distance-associated genes across different angular bins. **(G)** Expression of genes against intensity distance metric. Clustered by association type (e.g. negatively/positively associated).

**Supplementary Fig. 5.**
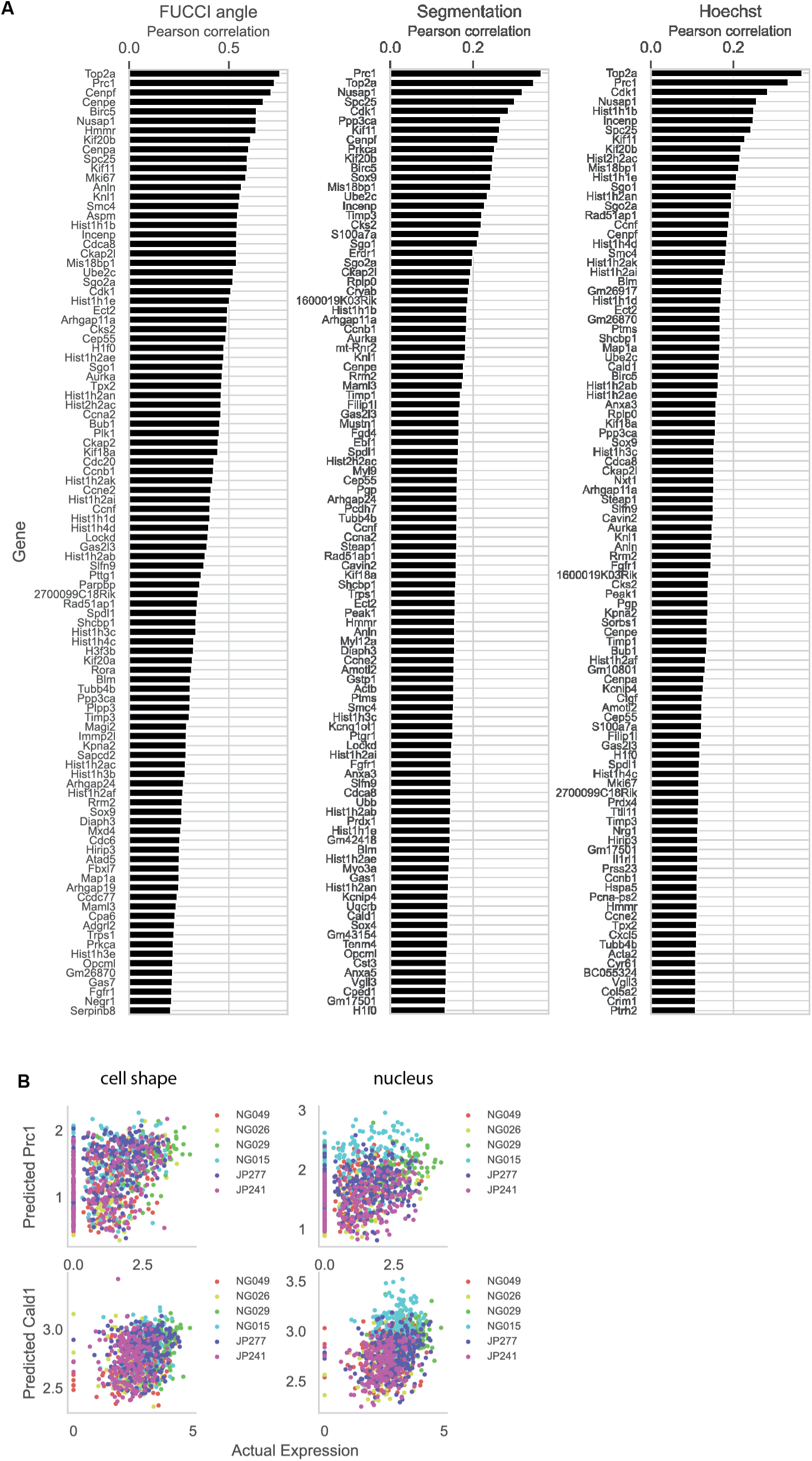
Machine learning models for predicting gene expression from cell morphology and angular features. **(A)** Top 100 genes by Pearson correlation for the cell shape, nucleus, and angular model. **(B)** Model predictions of the cell shape and nuclear model in relation to actual expression, colored by batch.

**Supplementary Fig. 6.**
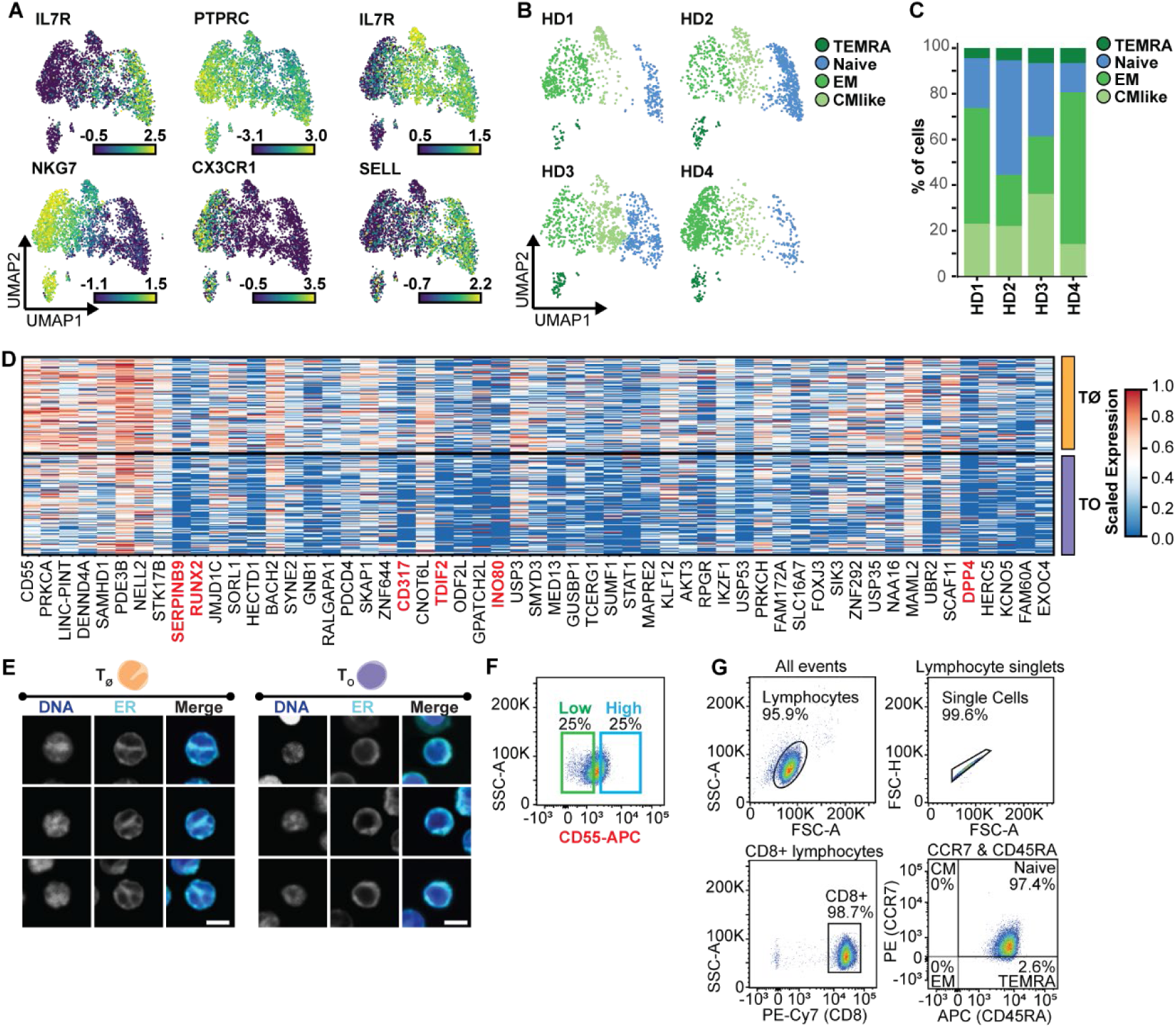
Differential gene expression across distinct naive CD8+ T cell morphologies identifies molecular targets for orthogonal validation. **(A-D)** Peripheral blood-derived CD8^+^ T cells (n = 3,969) across four donors were profiled with IRIS, pairing ER and nuclear morphology with scRNA-seq. **(A)** Expression of CD8^+^ T cell subset marker genes on the integrated UMAP. **(B)** Individual donors (HD) mapped onto the integrated CD8+ T cell embedding. **(C)** Proportion of each CD8^+^ T cell subset per donor. **(D)** Expanded heatmap (compared to Fig. 6H) of differentially expressed genes between TØ or TO naive CD8^+^ T cells on a per cell level. Significance determined by Wilcoxon test, Benjamini-Hochberg correction applied. Genes validated at the protein level are indicated in red. **(E)** Representative Hoechst and ER Bodipy stainings (single z-planes) used for confocal classification of T_Ø_ and T_O_ cells; ‘merge’ shows combined intensities. Scale bars, 5 μm. **(F)** FACS gating for low/high expression of CD55. **(G)** FACS gating strategy for naive CD8^+^ T cell sorting (CCR7 | T cell homing to lymphoid tissues, CD45RA | Activation history and maturation status).

## Material and Methods

### IRIS cell handling and isolation

#### Human blood

For experiments performed by the laboratory of Prof. Bart Deplancke buffy coats and whole blood were obtained from fully anonymized healthy donors provided by ‘Transfusion Interregionale’ (Biopole, 1066 Epalinges, Project-Nr. P_285).

For experiments performed by the laboratory of Prof. Berend Snijder, buffy coats were obtained from coded healthy donors provided by the Blutspende Zürich, under a study protocol approved by the Cantonal Ethics Committee, Zürich (KEK Zürich, BASEC-Nr 2019-01579).

#### Human PBMC and T cell enrichment

Peripheral blood (PB) was diluted 1:1 with DPBS (Thermo Fisher Scientific, #14190169). PB:DPBS was over-layered on Ficoll-Paque^TM^ PLUS (cytiva, #10351856) enabling purification of PBMCs by density centrifugation at 800 g for 30 min. The interphase containing PBMCs was washed once with DPBS and cell count was obtained with Neubauer (FAUST, #9161078) counting chamber with Trypan blue live–dead stain (Thermo Fisher Scientific, #T10283). When required, human CD8^+^ T cells were enriched from PBMCs using a column-based magnetic-activated cell sorting (MACS) Pan CD8 T cell Isolation negative selection kit (Miltenyi Biotec, #130-096-495) or for naive CD8+ T cells the Naive Pan T Cell negative selection isolation Kit (Miltenyi Biotec, #130-097-095) followed by CD8+ T Cell negative selection isolation kit (Miltenyi Biotec, #130-096-495), according to the manufacturer’s specifications.

#### Cell lines

Human HEK 293T cells (American Type Culture Collection (ATCC), #SD-3515), mouse NIH/3T3 (ATCC, #CRL-1658), Fluorescent ubiquitination-based cell cycle indicator Fucci(SA)5 mouse 3T3 cells (provided by Prof. Suter’s laboratory, EPFL Lausanne) were used.

#### Cell culture

HEK 293T and Fucci(SA)5 3T3 cells were cultured in DMEM (Thermo Fisher Scientific, #41966029) with 10% v/v FBS (Thermo Fisher Scientific, #A5670701) and 1% v/v penicillin–streptomycin (Thermo Fisher Scientific, #15140122). NIH/3T3 cells were cultured in DMEM Glutamax^TM^ (Thermo Fisher Scientific, #10566016) with 10% v/v FBS and 1% v/v penicillin–streptomycin. S2 cells were cultured in Schneider’s Drosophila Medium (Thermo Fisher Scientific, #21720024) with 10% v/v FBS and 1% v/v penicillin–streptomycin.

Cells were cultured to 70% confluency. Prior to use, cells were washed with DPBS, dissociated with 0.25% Trypsin-EDTA (Thermo Fisher Scientific, #25200056), washed with DPBS and counted with Neubauer counting chamber (FAUST, #9161078) with Trypan blue live-dead stain (Thermo Fisher Scientific, #T10283).

#### Live-cell organelle staining

Live cells were stained for 30 min at 37°C and 5% CO_2_. Endoplasmic reticulum was stained with 2 μM ER-Tracker^Tm^ (Bodipy) green (Thermo Fisher Scientific, #E34251) in DPBS (without Ca2^+^/Mg2^+^, Thermo Fisher Scientific, #14190169). F-actin was stained with 1 μM CellMask™ Deep Red Actin Tracking Stain (Thermo Fisher Scientific, # A57245) in DPBS (without Ca2^+^/Mg2^+^). Mitochondria were stained with 10 nM PKmito DEEP RED (Spirochrome, #SC055) in DPBS (without Ca2^+^/Mg2^+^). Lysosomes were stained with 1 μM SiR700 Lysosome (Spirochrome, #SC016) in DPBS (without Ca2^+^/Mg2^+^). CellTracker^TM^ dyes were used at 1 μM in DPBS. Nuclear stains were performed with 5 ng/mL Hoechst 34580 (Merck, #63493) in DPBS.

### IRIS experimental procedure

#### Microfluidic chip design and fabrication

The design of the microfluidic chip for deterministic co-encapsulation is presented in **Supplementary Fig. 2A**. The computer-aided design (CAD) files are available as Supplementary Data. Chips were designed using AutoCAD LT software (version 2020). The 5-inch chromium masks for both the control and flow layers were exposed in a VPG200 laser writer (Heidelberg instruments). Masks were developed using an HMR 900 mask processor (Hamatech). For the control layer, a 50 μm-thick SU8 photoresist layer was deposited with an LSM-200 spin coater (Sawatec), exposed on an MJB4 single side mask aligner (SÜSS MicroTec), and manually developed. The SU8 processing steps were carried out according to the manufacturer’s instructions for the 3050 series (Microchem). To fabricate the flow layer, a double resist coating process was performed to achieve a selective, locally variable channel cross-section. First, the rectangular cross-section channels were created by coating the wafers with a 50 μm-thick SU8 photoresist layer using an LSM-200 spin coater (Sawatec). The coated wafers were then exposed using an MJB4 single side mask aligner (SÜSS MicroTec) and manually developed. Next, to form the semi-circular cross-section valve areas, a 50 μm-thick AZ40XT (Microchem) positive photoresist was subsequently coated on the same wafer using the ACS200 system (Gen3, SÜSS MicroTec). The exposure step was performed on a MA6Gen3 mask aligner (SÜSS MicroTec) by carefully manually aligning the masks over the existing flow channels. Wafers were then developed on the ACS200 system. The developed master wafers were reflowed for 120 sec at 130°C on a hotplate until the valve areas appeared round under an inspection microscope. At this temperature, only the positive resist undergoes reflow, enabling the fabrication of devices with both rectangular and semi-circular channel cross-sections. The master-wafers were used as molds for polydimethylsiloxane (PDMS) chips after passivation with 1% silane dissolved in hydrofluoroether (HFE). The thick flow layer was obtained by mixing PDMS with curing agent at a ratio of 5:1 using a centrifugal mixer (Thinky), degassing for 15 min, and curing for 30 min at 80°C. The control layer was fabricated by spin coating the PDMS and curing agent at a ratio of 20:1 on the flow layer wafer at 1200 rpm. for 50 sec with a 9 sec ramp time, followed by baking at 80°C for 30 min. Cured PDMS was then detached from the flow layer mold and cut into individual chips, and inlet holes were punched with a 0.55 mm diameter biopsy punch (Darwin microfluidics, #PT-T983-05). The two PDMS layers were manually aligned and bonded at 80°C for at least 60 min. Assembled and cured PDMS chips were cut from the molds, and the control layer inlet holes were punched. Finally, chips were oxygen plasma activated (45 sec at ∼500 mTorr O_2_) and bonded to a surface-activated glass slide followed by incubation at 80°C for at least 2 h.

#### Microfluidic chip preparation

All Tygon tubing was connected to the inlet holes in the PDMS via 0.65/0.35 × 10 mm metal pins (UNIMED, #200.010-A) and nomenclature of inlets and outlets refers to **Supplementary Fig. 2A**. Quake valve inlets (i-v) were primed with water-filled Tygon tubing (Cole-Parmer, OD 0.06 inch, #06420-02) at 1.5 bar air pressure. The input channel for cell loading was connected via Tygon tubing to a 2 mL syringe (CODAN, #62.2611) connected via TE NEEDLE 23 GAUGE 1/4” (METCAL, #ZZ3260). The syringe, containing cells in suspension and an epoxy-coated magnet allowing for gentle stirring, (HKCM, #Z04×06Ep-N35) was placed in the combined cell mixer and cooler (Advanced Microfluidics, #CM1). The injection inlet for cell placement and stopping (2) was connected via Tygon tubing to a 200 μL Prot/Elec^Tm^ Tip (BIORAD, #12021138) containing 120 μL of 15% v/v Optiprep (Merck, #D1556) in DPBS. The inlet for the “cell barcode” (3) was connected via Tygon tubing to a 200 μL Prot/Elec^Tm^ Tip and loaded with 30 μL of the reverse transcription (RT) solution. The outlet for the RT solution exchange (4) and the liquid waste exit channel (8) were connected via Tygon tubing to a 7 mL waste reservoir. The oil inlet for droplet generation (5) and the oil inlet for droplet ejection (6) were connected via Tygon tubing to a pressurizable 5 mL syringe loaded with EvaGreen oil (BIORAD, #186-4006) connected via a TE NEEDLE 23 GAUGE 1/4”. The droplet exit channel was mounted with 3.5 cm of PEEK tubing 1/32” x 0.2 mm / 0.008” inner diameter (VICIJOUR, #JR-T-5708-M1). Subsequently, the chip was mounted in front of the vertically oriented objective and focused on the cell detection and droplet generation region (**Supplementary Fig. 2B**).

### IRIS instrument (hardware and software)

#### Microfluidic control system

All numeric valve annotation ((1)-(5), (i)-(v)) are displayed in **Supplementary Fig. 2A**. The pressures for microfluidic chip operation were digitally controlled with three types of valves: conventional proportional valves, pneumatic valves, and piezo proportional valves. All input pressures were first pre-regulated with four conventional proportional valves (Festo, #EAA-B-3-D9-F-V1-1R1). Four lines were used: 1. as pre-regulator for pneumatic valve bank 1, 2. as pre-regulator for pneumatic valves bank 2, 3. as pre-regulator for the piezo proportional valves, 4. the pressure for the oil flush reservoir (6).

The input lines of the piezoelectric proportional valve and pneumatic valve bank 1 were buffered (Festo, CRVZS0,1).

The pneumatic valves (Festo, MHA1-M1H-3/2O-0,6-HC) were separated into two banks: bank 1 of 4 valves that controlled the cell valve (i), the sort valve (v), the waste valve (iv), and the biochemistry wash outlet valve (iii). Bank 2 comprised one valve that controlled the leak valve (ii). Bank 1 was set to 1.4 bar pressure, bank 2 to approximately 900 mbar and was fine-adjusted when necessary.

The piezoelectric proportional valves (Festo, VEMP-BS-3-13-D19-F-28T1) were supplied with 1.7 bar of pressure from one conventional proportional valve. The piezo valves were controlled with a piezo control board (Octanis instruments, High Voltage Piezo Driver) and pressures measured with a pressure sensor (Honeywell, ABPMANV030PGAA5). Four piezoelectric valves were connected to the cell reservoir (1), the leak-flow inlet (2), the biochemistry inlet (3), and the oil inlet (5).

#### Microscopy system

The microscope comprised of an objective (Nikon, S Fluor 40x NA0.9), a camera for imaging (PCO, Edge 4.2 USB) with tube lens (Nikon, 58-520), and a multicolour laser source (Hubner Photonics, C-Flex C4, fiber couple Kineflex APC, 405 nm 150 mW, 488 nm 200 mW, 561 nm 100 mW, 638 nm 180 mW) for illumination. The excitation laser beam was collimated (Thorlabs, AC254-050-A-ML), the beam expanded (Thorlabs, GBE02-A), the excitation laser was focused with a 200 mm lens (Thorlabs, LA1708-A) on the back focal plane of the objective. The laser was coupled with the imaging camera over a quadband filter set (Chroma, 89402ET), comprising a dichroic, an excitation filter, and an emission filter. The detection camera light was filtered (Thorlabs, DMLP805) and coupled over a dichroic (Thorlabs, DMSP805R). A combination of a 20 mm lens (Thorlabs, ACL2520U-B) and a 50 mm lens (Thorlabs, LA1131-AB) was used as tube lens and beam de-expander.

The wide-field condenser located on the opposite side of the objective comprised a white LED (Thorlabs, MNWHL4) for imaging illumination with a 520 nm filter (Thorlabs, FBH520-40), and a near infrared LED (Thorlabs, M850L3) for detection illumination, both driven by a controllable driver (Thorlabs, LEDD1B), both equipped with a collector lens (Thorlabs, ACL2520U-B), a field diaphragm (Thorlabs, CP20S), and their beams combined over a dichroic (Thorlabs, DMSP805R). The dichroic was followed by a field lens (Thorlabs, AC254-050-AB-ML) and the condenser lens (Thorlabs, ACL25416U). An aperture diaphragm was only temporarily introduced for alignment and removed after.

The chip was fixed on a piezo positioner (Piezo System Jena, MICI 200 SG) controlled by piezo amplifier (Piezo System Jena, 12V40 SG OEM) that moved the chip forward and backward for focusing.

#### Machine-vision hardware and software

The system was controlled by an embedded system (AVNET, Ultra96-V2) running Petalinux (version 5.4.0) and operated by custom software adapted from previously developed software (https://github.com/DeplanckeLab/DisCo_source). Briefly, images were captured by the detection camera (Daheng Imaging, MER-031-860U3M-L) connected via USB using the Daheng Imaging Galaxy Linux Arm SDK (version 1.3.2). For cell detection, images were processed with the OpenCV computer vision library (version 3.4.3). Regions of interest (ROI)s were extracted from the raw images by cropping, the cropped images were denoised by Gaussian blurring, and two images 10 ms apart subtracted, the resulting image binarized by thresholding, and holes filled by dilation. In the processed binaries, cells were detected by contour detection using the *findContours* function, and contours thresholded by area and circularity. Imaging data from the PCO Edge camera was captured on a separate computer connected to the PCO camera via USB and utilizing the pco Python API (version 0.1.3) for image capture.

Cell positioning was achieved in two steps. Cells were first detected in the first region of interest (ROI1, **Fig. 2A**). Upon cell detection, the cell inlet ‘Quake’-valve was closed. Next liquid was flown from the leak inlet while cell detection was run on the second ROI (ROI2, **Fig. 2A**). Upon detection (and thus cell positioning), the leak flow was stopped. Subsequent to cell stopping, a plug was formed via 150 msec over-pressurization of the leak-flow ‘Quake’-valve from the ‘RT’ solution inlet (**Video 1**). The plug containing the cell in 2.5 nL of running buffer and 2.5 nL of RT solution (see section ‘Library preparation’), was sheared by over-pressuring the leak-flow ‘Quake’-valve from the dropletting oil inlet and ejected into the sort outlet (**Video 2**). Each droplet containing one cell was ejected from the chip with 3.5 μL of dropletting oil (BioRad, #186-4006) into one well of 96-well plate (StarLab, #4ti-0760). The 96-well plate was automatically moved row-by-row on a per well basis via an XY-stage to capture the ejected droplet.

### IRIS molecular profiling

#### Generation of evaporation protection solution

Evaporation protection solution serves as an evaporation protection for the reaction droplets containing the cells and RT solution, during the droplet consortia assembly. 96-well plates were loaded with 1 μL of 1x RT buffer (Thermo Fisher Scientific, #EP0753) containing 7.5% v/v Optiprep (Thermo Fisher Scientific, #D1556). Then, 5 μL of dropletting oil was added to each well. The plate was sealed (Merck, #CLS6570-100EA) and centrifuged at 1000 g for 1 min. The sealed 96-well plate was vortexed for 10 s on a Vortex Genie 2 (Scientific Industries, #SI-0236) at speed ‘10’. The vortexed 96-well plate was centrifuged at 1000 g for 1 min and then used to assemble droplet consortia.

#### Droplet consortia assembly

Droplet consortia were assembled by iteratively collecting 96 cells in droplets on chip with the same RT solution with one known cellcode primer, then exchanging the RT solution with a new cellcode primer, and then repeating the cell encapsulation for 96 cells. In brief, 96 cells were successively captured in droplets, individually ejected from the chip and deposited on a per-well basis in a 96-well plate containing the evaporation protection solution. Subsequently, the RT solution inlet was washed two times with 125 μL of H_2_O (Thermo Fisher Scientific, #10977035). Then, 30 μL of cellcode primer-containing RT solution was added to the protein/electrophoresis tip (Bio-Rad, #12021138) connected to the RT inlet Tygon tubing. Next, the second set of 96 cells was individually encapsulated, ejected from the chip and deposited in the wells already containing one droplet with one cell. The process was repeated for up to 13 times to assemble ‘droplet consortia’ on a per well basis with the equivalent of up to 13 individual cells for single cell molecular profiling per well. Between one to eight 96-well plates were used per experiment.

#### cDNA generation from assembled droplet consortia

The generation of cDNA amenable for scRNA-seq consists of 1) the droplet-based reverse transcription that molecularly barcodes each cell, 2) the overhang-PCR amplifying the single-stranded cDNA and performing molecular barcoding per well, and 3) the amplification of well-coded droplet consortia-derived cDNA on a per 96-well plate basis.

All reverse transcription solutions used for assembling the consortia were of identical making and only differed in the reverse transcription primer used to capture the poly-adenylated mRNA (**Fig. 2A, Supplementary_Table_PRIMER**). The RT solution was composed of 1.75x RT buffer, 15% v/v Optiprep, 1% v/v Triton-X 100 (Thermo Fisher Scientific, #327371000), 2 mM dNTP (Merck, #DNTP100-1KT), 3 mM dCTP (Thermo Fisher Scientific, #10217016), 0.01M Tris-HCl pH 8.0 (Thermo Fisher Scientific, #15568025), 6.8 mM DTT (Merck, #43815), 0.01 mM MgCl2 (Merck, #M1028), 2 U/μL RNAseOUT (Thermo Fisher Scientific, #10777019), 20 U/μL, RT Maxima Enzyme (Thermo Fisher Scientific, #EP0753), 1.6 μM RT-TSO primer and 0.5 μM RT-cellcode primer (**Supplementary_Table_PRIMER**). Reverse transcription was carried out within the individual droplets of the assembled droplet consortia at 46°C for 60 min followed by 5 min at 85°C.

Each 96-well plate was processed individually. Prior to performing the wellcoding PCR, droplets were merged by applying an alternating electric field of 10kV at 50 Hz. Subsequently, to each well, 9 μL PCR reaction mix was added composed of 1.1x Terra^Tm^ Direct Polymerase Mix (Takara, #639271) containing 0.16 μM wellcoding primer and 0.32 TSO-PCR primer. The 96-well plate was sealed, centrifuged at 1000 g for 1 min and PCR carried out as follows: (3 min 98°C) --> 5x cycles [(20 sec 98°C)(45 sec 57°C)(3 min 72°C)] --> (10 min 72°C) --> (hold 4°C). Liquids of all wells were pooled in a 15 mL Falcon, and centrifuged at 1000 g. Then, 4.2 mL of PB binding buffer (QIAGEN, #19066) was added, the solution applied to a QIAquick Spin Column (QIAGEN, #28115) and the cDNA captured on the column using the QIAvac 24 Plus (QIAGEN, #19413). cDNA was recovered according to the manufacturer’s instructions of the QIAquick PCR Purification Kit (QIAGEN, #28104) eluting the cDNA in 100 μL H_2_O. Bead-based size-selection was performed at 0.8x bead-to-elution ratio according to the manufacturer’s instructions using CleanNGS (CleanNA, #CNGS-0500). cDNA was eluted in 50 μL H_2_O. PCR reaction mix was added, composed of 2x Terra^Tm^ Direct Polymerase Mix containing 0.16 μM P5 and 0.32 μM TSO-PCR primer (**Supplementary_Table_PRIMER**). PCR was carried out as follows: (3 min 98°C) --> ‘N’x cycles [(15 sec 98°C)(30 sec 60°C)(4 min 72°C)] --> (10 min 72°C) --> (hold 4°C). ‘N’ corresponds to the number of PCR cycles that varies along cell type. For example, PBMCs were amplified for 10 PCR cycles for 480 cells/plate whereas cell lines were typically amplified for 7 to 8 PCR cycles for 480 cells/plate. Left-side bead-based size-selection was performed at 0.8x bead-to-elution ratio according to the manufacturer’s instructions using CleanNGS (CleanNA, #CNGS-0500). Fragment Analyzer (Agilent, DNF-474-0500 kit) and the Qubit High Sensitivity kit (Invitrogen, Q33231) were used for cDNA quality control and quantification.

#### Tagmentation

cDNA processing for sequence library preparation was carried out according to Picelli *et al*^58^. Each 96-well plate was indexed individually. Tn5 was produced in-house (EPFL, Protein Production and Structure Core Facility) and B/Read2/P7|/B/Read2/P7 loaded (**Supplementary_Table_PRIMER**). Tagmentation was carried out with 2 ng of cDNA for 5 min at 55°C. To stop tagmentation, SDS was used at 0.2% (Merck, #71736). For library amplification, the Kapa HiFi kit with dNTPs (Roche, KK2102) was used according to the manufacturer’s instructions containing 0.3 µM P5 and 0.3 µM PC7xx primer (**Supplementary_Table_PRIMER**). PCR was carried out as follows: (3 min 72°C) --> (30 sec 98°C) --> 14x cycles [(10 sec 98°C)(30 sec 63°C)(60 sec 72°C)] --> (5 min 72°C) --> (hold 4°C). Two subsequent left-side bead-based size selections at 0.7x and one left & right-side bead-based size selection 0.70x-0.50x were carried out using CleanNGS. Fragment Analyzer and the Qubit High Sensitivity kit were used for cDNA quality control and quantification.

#### Sequencing strategy and demultiplexing

Sequencing libraries were sequenced either on a NextSeq 500 (Illumina) with i5-10bp, i7-10bp, Read1-16bp and Read2-59bp or on an AVITI (Element Bioscience) with i5-10bp, i7-10bp, Read1-16bp and Read2-129bp. NextSeq 500-generated BCL files were demultiplexed using *bclconvert* (Illumina) according to the manufacturer’s instructions (https://emea.support.illumina.com/sequencing/sequencing_software/bcl-convert.html) and AVITI-generated BCL files were demultiplexed using *bases2fastq* (Element Biosciences) according to the manufacturer’s instructions (https://docs.elembio.io/docs/bases2fastq/), both with default settings. Demultiplexing was performed based on the i5 and i7 indexing sequences providing demultiplexed fastq files for Read1 and Read2 on a per plate per well basis (**Supplementary_Table_PRIMER**). The first 7 bp of Read1 contain the cellcode and the remainder 9 bp contain the UMI information. Read2 contains the transcript information.

#### Sequence data analysis

Alignment of reads was performed on a per plate per well basis. Reads were aligned to the human (hg38), mouse (GRCm38), fly (dmelr6.37), or combined reference genomes hg38&GRCm38 or hg38&GRCm38&dmelr6.37, depending on the origin of the cellular input material. Alignment was performed using *STARsolo* (version 2.7.9.a)^59^ demultiplexing the paired-end Read1 and Read2 to the respective cell codes and transcript count utilized for the given experiment. The resulting count matrices per well were compiled per experiment into a count matrix per experiment. Each cell is identifiable by a combination of four unique identifiers being the experiment ID (expID), the plate code (PC, corresponding to the i7), the well code (WC, corresponding to the i5) and the cell code (CC, corresponding to the first 7bp of the Read1). Each cell is therefore fully traceable throughout the experiment by its unique Full-Cell-ID structured as ‘expID_PC_WC_CC’ (e.g., PBMC1_PC1_WC42_CC1).

#### Image data pre-processing and feature extraction

Brightfield images of the entire stopping zone (512 x 2048 pixels) were acquired after each successful cell detection event. For each event, image stacks across multiple focal planes were collected. These image stacks were processed using the YOLO object detection algorithm (v8m)^60^, which was retrained on our dataset for cell detection and localization. YOLO provides both the coordinates of bounding boxes surrounding each detected cell and the index of the focal plane where the cell is in focus. These coordinates were subsequently used to generate 250 x 250 pixel image crops, cantered around each cell, for each available fluorescent channel acquired.

For cell segmentation, the 250 x 250 brightfield crops were processed using DeepLab v3^61^, a semantic segmentation architecture. The network was retrained on several thousand brightfield images from our dataset to achieve accurate segmentation of single-cell outlines. The output of DeepLab v3 is a binary mask highlighting pixels belonging to the cell from background ones. This segmentation mask was used to compute cell-level quantitative image features, taken at the focal plane in focus as determined by YOLO. The cell size was calculated as the number of nonzero pixels in the binary segmentation mask and converted to µm² using the image resolution. The circularity was computed from the segmentation mask contour. Fluorescent intensities were extracted for each available channel, as the mean and max pixel intensity values within the segmented cell region.

### FUCCI cell cycle analysis

#### scRNA-seq data analysis (FUCCI dataset)

Analyses were performed using Scanpy^62^ (v1.10.2) following standard practices. In brief, cells were quality filtered to retain high quality cells and genes (min_genes ≥ 1000, pct_counts_mt < 15%), genes selected (min_cells ≥ 0.5%), counts normalized and log-transformed, and dimensionality reduction (PCA, UMAP; n_pcs=60) was performed. Different experiments were integrated using Harmony^63^ (v0.0.10).

#### Intensity and angular score calculation

Intensities were calculated by masking the image and calculating the average intensity over the masked image. Based on the mean intensities, a continuous angular score ranging from 0 to 2π was computed for each cell, as previously described^35^. The log-transformed (log1p) FUCCI intensities were projected onto the orthonormal basis defined by the first two principal components PC1 and PC2. The angular score was then computed using the two-argument arctangent function as θ=atan2 (PC2, PC1), yielding continuous angular values in (0, 2π).

#### Differential gene expression analysis across angular cell cycle bins

Based on the continuous ‘cell cycle angle’ score, cells were grouped into 50 discrete angular cell cycle bins of equal angle (‘window’). Differential gene expression analysis was performed for each window against all other groups using the function scanpy.tl.rank_genes_groups() (method=’wilcoxon’) on the subset of top 2000 highly variable genes. DEGs were filtered based on significant adjusted p-value (≤ 0.05).

#### Visualization of DEGs across angular cell cycle bins

Angular bin specific genes as shown in **Fig. 4D** were chosen by selecting the top 3 up-regulated DEGs per angular-bin without repetition across bins. DEGs from angular bins were compared with the Tirosh et. al.^37^ gene list and GO cell-cycle genes by downloading the full list of genes associated to GP:BP “cell cycle” (ID: GO:0007049) and the list of Tirosh et. al. ^37^ genes. On the set of genes not overlapping with either the Tirosh et. al. ^37^ or GO cell cycle gene lists, gene set enrichment analysis was performed using EnrichR web tool^64^ on databases “GO_Biological_Process_2021”, “GO_Cellular_Component_2021”, “Reactome_Pathways_2024”, “TISSUES_Curated_2025”.

#### Cell-cycle annotation with Seurat and tricycle

Seurat cell-cycle annotation was performed using Scanpy’s implementation^65^, with (*scanpy.tl.score_genes_cell_cycle()*) using the cell cycle reference panel of genes curated by Tirosh et al.^37^.

Tricycle cell-cycle annotation^39^ (v1.10.0) was performed according to recommended practices: cells were projected onto the tricycle pre-learned cell cycle space (*scater::plotReducedDim()*) and the cell-cycle position estimated using function *estimate_cycle_position()*. For plotting purposes, the continuous position score was discretized to cell-cycle phases according to thresholds suggested by the authors (0.5pi: S stage start; pi: G2M stage start; 1.5π: middle of M stage; 1.75π-0.25π: G1/G0 stage). Continuous and discrete annotations for single cells were exported and embedded in the scanpy adata object. Cell-cycle annotation methods were compared by assessing the percentage of cells classified to the same phases.

#### Transcription factor target gene enrichment analysis

Transcription factor target gene enrichment analysis on angular-bin DEGs was performed with gseapy library’s^66^ (v1.1.3) enrichr() function on target gene sets: “TRRUST_Transcription_Factors_2019” and “ENCODE_and_ChEA_Consensus_TFs_from_ChIP-X”. Significantly enriched TFs (adjusted p-value ≤ 0.05) were retained. For each angular-bin, top enriched TFs according to significance were selected and plotted.

#### Dream complex activity

Dream Complex activity was inferred using a publicly available Dream target genes list^42^. The list was filtered to retain genes being differentially expressed in our setting (firstly: across cell-cycle angular score, secondly: along ‘intensity distance’ metric) and used to compute a “Dream signature score” across single cells using *scanpy.tl.score_genes()*.

#### Differential gene expression analysis along ‘intensity distance’ metric

The ‘intensity distance’ metric for each cell from the center point was determined by calculating the Euclidean distance of each cell from the center and normalized using min-max normalization. To identify genes whose expression varied as a function of the intensity distance metric, a linear regression between gene expression and the intensity distance metric, using *statsmodels* (*statsmodels.OLS.fit()*),), was performed: gene-wise, cyclically for each angular bin, and on HVGs. Resulting p-values were corrected for multiple testing to adjusted p-values with the Benjamini-Hochberg (BH) method. Genes with a significant adjusted p-value (≤ 0.05) were considered differentially expressed and their positive or negative regulation with respect to intensity distance metric determined based on the linear regression coefficient. The matrix of DEGs x cells sorted based on intensity distance metric was clustered using *seaborn.clustermap()* to detect shared patterns of expression. Specific clusters of negatively-associated DEGs were selected for plotting.

#### CNN model architecture and training

We trained ResNet18^44^ convolutional neural networks (CNNs) with regression outputs to predict expression of the top 2,000 HVGs from two input types: nuclear morphology (Hoechst) and whole-cell shape masks from brightfield images. The output layers of the ResNet18 were replaced with a fully connected layer with one output unit per gene. The nuclear model input training data was augmented by applying random horizontal flips with 50% probability and random ±90° rotations with 50% probability. Additionally, random changes in brightness and contrast of the input image (brightness factor range: −0.01 to 0.01, contrast factor range: −0.01 to 0.01, probability of applying the transformation: 20%) were applied and Gaussian noise (standard deviation range: 0.005 to 0.01, probability of applying the transformation: 100%) added. For the cell shape model, augmentations were restricted to random ±90° rotations with 50% probability. The raw counts of the gene expression data were normalized to the median library size (*scanpy.pp.normalize_total* with default arguments), followed by a log transformation *scanpy.pp.log1p*. The cells in the dataset were randomly split into train (60%), validation (20%), and test (20%). The models were trained to minimize the mean squared error, yielding the conditional mean of gene expression values given the input images. Training was performed for up to 50 epochs with batch size 256, and early stopping on the validation loss was applied with a patience of 10 epochs. Models were optimized using the Adam optimizer (learning rate 0.001 and weight decay 0.0001) and the learning rate was decayed every 10 epochs with a multiplicative factor of 0.1.

#### CNN and MLP Model evaluation

To eliminate the influence of technical variation on the evaluation metrics, the model performance was evaluated within each experimental batch by computing gene-wise Pearson correlations between predicted and observed expression. We averaged the per-gene correlations across experiments and combined the Pearson p-values with Fisher’s method. To increase the robustness of the metrics, three independent models were trained with random train/validation/test splits. The per-gene performances were calculated by averaging the test scores across the three independent runs. The false discovery rate was controlled at α=0.05 using the Benjamini-Hochberg procedure.

#### Multi-layer perceptron (MLP) model

The multilayer perceptron (MLP) was trained on the continuous cell cycle angular score. An MLP was chosen due to the nonlinear, circular nature of the cell cycle. The MLP consisted of 1 hidden layer (32 units) with a rectified linear unit (ReLU) activation and one output unit per gene. The model was trained with the mean squared error loss, a batch size of 256 cells, and the Adam optimizer (learning rate 0.01 with no weight decay). Training was run for up to 500 epochs with early stopping on the validation loss (patience 2). Gene expression preprocessing and data splits matched those used for the CNN models.

#### Angular bin DE-genes vs CNN performance

For each angular bin the predictive CNN performance (Pearson correlation coefficient on test data) was averaged over all genes contained in the respective bin.

#### PCA of predictable genes

Principal component analysis (*PCA(), pca.fit_transform()*) was performed on the expression matrix of all genes with significantly (p < 0.01) associated with all models (cell shape, nucleus, angular). Genes distribution according to the first two principal components was visualized.

#### Relative performance calculation and GSEA

All genes with a significant association (p < 0.01) for both morphology models (cell shape and nucleus model) were selected. For each model (cell shape, nucleus, angular) all Pearson correlation values were normalized by the maximum Pearson correlation, respectively. For each gene a relative performance value for the cell shape and nucleus model was computed by dividing the respective absolute normalized Pearson correlation value by the absolute normalized correlation value of the angular model. For gene set enrichment analysis (GSEA), genes with a 2-fold higher relative performance than the relative angular performance were selected for both cellular shape and nucleus model, respectively. Genes obtained this way were gene set enriched with gseapy (V1.1.4) and the ‘Reactome 2022’ gene set.

#### Linear regression of circularity and cell area

Calculation and models were calculated/fitted for the cells in the test dataset for the morphology and an additional filter step was applied to remove small cell contours of < 5000 pixel in size. Measurements were computed with OpenCV Python (V4.10.0.84). Cell segmentation masks of the in-focus plane were used to calculate cellular contours (findContours). Area and perimeter of the cells were computed from the contours with contourArea and arcLength functions. For each gene, a linear regression model was fitted using scikit-learn (V1.7.2) and the Pearson correlation of predictions from the regression models with the predictions from the morphology models calculated.

#### Timelapse microscopy and data analysis

3T3 FUCCI cells were cultured at 37°C and 5% CO2 and imaged every 30 min for 17 h on a Leica DMi8 inverted thunder imager with a HC PL FLUOTAR 10x/0.32 dry objective. 100 cells from the same field of view were manually tracked and maximum intensities calculated. Each intensity measurement was corrected for global intensity variation across the field of view by manually fitting a compensation function. The background intensity of each channel was subtracted by calculating the global minimum intensity. The angular score was calculated as described with the only alteration that the log10 intensities were scaled with the scikit-learn PowerTransformer function before applying PCA. To calculate the angular difference the phase was unwrapped for each track (all measurements of one cell) by numpy’s unwrap function, and the minimum unwrapped value subtracted from the maximum value. A regression model was fitted to the angular difference and maximum AzaleaB5-hCdt intensites using the statsmodel libraries ordinary least squares (OLS) function.

### CD8^+^ T cell validation

#### scRNA-seq data analysis (PBMCs dataset)

Analyses were performed using Scanpy (v1.10.2 – 10.1186/s13059-017-1382-0) following standard practices. Data was filtered to retain high quality cells and genes (min_cells ≥ 0.2%, min_genes ≥ 1000, pct_counts_mt < 15%), normalized and log-transformed, and dimensionality reduction (PCA, UMAP; n_pcs=27) was performed. Cell cycle scoring was performed using Scanpy’s implementation of the Seurat approach^65^, with (scanpy.tl.score_genes_cell_cycle()) using the cell cycle reference panel of genes curated by Tirosh et al.^37^. Different experiments were integrated using Harmony (v0.0.10 – 10.1038/s41592-019-0619-0). Cells were grouped into clusters (*sc.tl.leiden()*, resolution=0.15) annotated based on marker gene expression. Differential gene expression across clusters was performed with sc.tl.rank_genes_groups(method=’wilcoxon’) and default thresholds. For downstream analyses, two clusters (’HLA-DRA+’, ‘unknown1’) were ignored as not T cells and thus irrelevant to the study. An additional clustering step was performed on this final UMAP (resolution=0.3).

#### Analyses on ‘stripy’ and conventional T cells

Stripy and conventional T cell annotations were transferred from single cell image annotations obtained through manual curation of cell morphologies based on staining with ER-Tracker^Tm^ (Bodipy) green (Thermo Fisher Scientific, #E34251) for organization of the ER around the nucleus, being TO, and cells with the ER being localized centrally in an elongated shape, being TØ. Differential gene expression analysis between stripy and conventional cells was performed for each T cell cluster (*<u>sc.tl</u>.rank_genes_groups()*), significant DEGs were filtered based on adjusted p-value and log fold-change thresholds (≤ 0.05 and ≥ 0.5, respectively). GO Biological Process Gene set enrichment analysis on up-regulated genes in CD8 Naive Stripy cells vs. CD8 Naive conventional cells was performed.

#### Human peripheral blood mononuclear and T cell enrichment

Peripheral blood was diluted 1:1 PBS (Gibco; ThermoFisher Scientific, Waltham, MA), before addition of Lymphoprep (Stemcell Technologies, Vancouver, Canada) and isolation of PBMCs by density centrifugation at 800 g for 30 min. PBMCs were washed twice with PBS and cells were counted (Countess II, ThermoFisher Scientific). Naive CD8 T cells were enriched from PBMCs using sequential column-based magnetic activated cell sorting (MACS) isolation using the Pan naive T cell Isolation Kit (Miltenyi Biotec, #130-097-095) followed by the Pan CD8 T cell Isolation Kit (Miltenyi Biotec, #130-096-495), according to the manufacturer’s specifications. Cells were resuspended in RPMI (Thermo Fisher Scientific, #72400021) supplemented with 10% FBS and 10% DMSO (Merck, #41639), and slow frozen to −80°C using a Mr Frosty^Tm^ freezing container (Thermo Fisher Scientific, #5100-0001).

#### Fluorescence-activated cell sorting

MACS-enriched human T cells were resuspended to 5×10^7^ cells/mL in DPBS supplemented with 0.2% w/v Bovine Albumin Serum (Merck, #A7906) containing 0.5 mM EDTA (Thermo Fisher Scientific, #AM9260G) and stained with respective antibodies for 20 min at 4°C (**Supplementary_Table_Antibodies**). Cell sorting was performed with an Aria II flow cytometer (BD Biosciences). FACS data were analyzed using FlowJo software (FlowJo Enterprise, version 10.0.9). For cell culture, T cells were collected in 10% FBS in RPMI and concentrated to approximately 400,000 cells/mL prior to seeding into clear-bottom PhenoPlate 384-well plates (Revvity, #6007460). Cells were plated in at least triplicate wells, and shortly spun down at 100g to ensure cells rested on the bottom of the wells. Cells were incubated for 30 min at 37°C with 5% CO_2_.

#### T cell staining for automated microscopy

T cells were fixed and permeabilized for 20 min at room temperature with 1% (w/v) formaldehyde (Merck, #104002), 0.05% v/v Triton X-100 before blocking at 4°C with PBS supplemented with 5% BSA. For staining of T cell architecture, conjugated anti-CD3 (**Supplementary_Table_Antibodies**) antibodies were diluted in DPBS containing 10 μM DAPI (Merck, #D9542) and added to blocked cells in the dark at room temperature for one hour. Cell solutions were aspirated, and DPBS added to cells prior to confocal imaging. Automated fluorescence microscopy was performed with an Opera Phenix automated spinning-disk confocal microscope (PerkinElmer). Human-derived antibody-stained T cells were imaged with a 20x 0.4NA air objective with 5×5 non-overlapping images capturing the entirety of each well. The acquired raw .tiff images were automatically transferred from the microscope for subsequent analysis.

#### Convolutional neural networks for automated classification of T cell architecture

T cell architecture was classified as previously described, using the T_NET_α network^46^. T_NET_α was trained to classify fixed and permeabilized human T cells as TØ or T_O_ cells via DAPI. Brightfield and CD3 48 x 48 pixel (14.4µm x 14.4µm) single cell crops were imaged at 20x 0.4NA. The training set comprised 12253 T_Ø_, 14396 T_O_ and 8437 T_Ρ_ cells, derived from primary human samples. Testing was performed on 1535 TØ and 1795 TO.

#### High-resolution fluorescence microscopy

T cells were fixed and permeabilized for 20 min at room temperature with 1% w/v formaldehyde, 0.05% (v/v) Triton X-100 before blocking at 4°C with DPBS supplemented with 5% FBS, using μ-Slide VI imaging slides (Ibidi, #80606). Primary unconjugated antibodies were diluted in PBS and used to stain fixed and permeabilized cells overnight at 4°C at the indicated dilutions (**Supplementary_Table_Antibodies**). After staining, cells were subsequently washed twice with DPBS. Conjugated primary and secondary antibodies were diluted in DPBS containing 10 μM DAPI and added to cells in the dark at room temperature for one hour. Cells were washed once with DPBS. Multi-channel fluorescence imaging was performed using a Nipkow spinning disk microscope coupled with a W1-T2 Confocal Scanner Unit (Yokogawa) and a 100x 1.45 CFI Plan Apo Oil objective. Images comprised 51 x 500 nm z-stacks, which were processed using ImageJ^67^ and MATLAB.

All T cells were cropped via ImageJ, and segmented using a custom script in MATLAB. For each cell, the sum projections were calculated for each fluorescent marker corresponding to the respective intracellular and the anti-CD3 antibody stain, based on the segmentation mask. The sum projections for each cell were normalized by the mean sum projections of all cells analysed for each marker and donor. T cell architecture was manually annotated by assessing the CD3 and DAPI organization for each cell. Only cells whose architecture was clear when visualizing max projections, and scanning through individual z-stacks, were considered for further analysis.

#### Statistics

Statistical analyses for validations on TØ and TO cells were performed using two-sided Student’s t-tests to assess differences between groups. Data are presented as mean ± standard deviation, and p-values are reported directly in the figures and legends.

## Acknowledgements

The authors thank the former members of the Deplancke laboratory (EPFL) for their support: D. Alpern and R. Dainese for constructive discussions. Furthermore, C. Albert, R. Rysman and M. Gevers from Advance Microfluidics for the custom manufacture of the cooled air-pressure-driven CellMixer, Duygu Koldere Vilain for illustrations and the EPFL CMi, GECF, BIOP, FCCF, Histology Core Facility, SCITAS, and UNIL VITAL-IT for device fabrication, sequencing, imaging, sorting, histology, and computational support, respectively. This research was supported by the Swiss National Science Foundation Advanced Grant (TMAG-3_209335) and a Precision Health and Related Technologies (PHRT-573) grant (to B.D.), the Swiss National Science Foundation SPARK initiative (CRSK-3_190627, to J.P.), the EuroTech PostDoc Programme co-funded by the European Commission under its framework program Horizon 2020 (754462, to J.P.), the SNSF MD-PhD program (323530_214540, to T.F.) and the Martinez Family Trust Fellowship in Computational Cancer Biology (to C.L.).

## Author information

These authors are equally contributing shared-first: J. J. Bues, J. Pezoldt, C. Lambert.

These authors are equally contributing shared-second: B. D. Hale, E. Bugani

J.B., J.P. and B.D. designed the study.

J.B., C.L., J.P. and B.D. wrote the manuscript.

C.L. and J.B. designed and fabricated the microfluidic chips.

JP., C.L., J.B. and A.H. optimized the on-chip microfluidic fluid workflow.

J.B., M.K. and A.H. developed the optical train.

J.V. assisted to the optimization of the microscope.

J.B. developed the integration for IRIS.

J.P., N.G., R.A., K.E., A.O. and S.S. established the scRNA-seq biochemistry.

J.P., N.G. and C.L. benchmarked the system and performed all single-cell RNA-seq experiments.

J.P., C.W. and V.G. established the scRNA-seq processing pipeline.

B.H. and S.L. performed the T cell experiments.

B.H. analyzed the T cell experiments.

J.B., C.L. and J.L. performed the FUCCI time-course experiments.

J.B. analyzed the FUCCI time-course experiments.

J.P., E.B., J.B. and A.J.C.O. performed data analysis related to benchmarking and T cell experiments scRNA-seq.

J.B., E.B., C.L. and M.A. performed data analysis related to FUCCI scRNA-seq.

J.B., C.L. and T.F. established the image pre-processing pipeline.

J.B., C.L. and R.T. performed image data analysis.

J.P. and N.G. performed cell culture and T cell isolation procedures.

D.P. supervised aspects of the optical train development.

W.K. supervised aspects of the time-course FUCCI experiments.

M.B. supervised the image-analysis and machine-learning approaches and reviewed the manuscript.

B.S. supervised the validation of the T cell experiments and reviewed the manuscript.

## Ethics declarations

B.D., J.B. and J.P. have filed a patent application for the deterministic co-encapsulation system (patent no. US11872559B2 and US20240401026A1). B.D. is co-founder of and has equity interest in Alithea Genomics. All other authors have no competing interests

